# A neuro-immune axis of transcriptomic dysregulation within the subgenual anterior cingulate cortex in schizophrenia

**DOI:** 10.1101/2025.02.14.638357

**Authors:** Rachel L. Smith, Agoston Mihalik, Nirmala Akula, Pavan K. Auluck, Stefano Marenco, Armin Raznahan, Petra E. Vértes, Francis J. McMahon

## Abstract

Many genes are linked to psychiatric disorders, but genome-wide association studies (GWAS) and differential gene expression (DGE) analyses in post-mortem brain tissue often implicate distinct gene sets. This disconnect impedes therapeutic development, which relies on integrating genetic and genomic insights. We address this issue using a novel multivariate technique that reduces DGE bias by leveraging gene co-expression networks and controlling for confounds such as drug exposure. Deep RNA sequencing was performed in bulk post-mortem sgACC from individuals with bipolar disorder (BD; N=35), major depression (MDD; N=51), schizophrenia (SCZ; N=44), and controls (N=55). Toxicology data dimensionality was reduced using multiple correspondence analysis; case-control gene expression was then analyzed using 1) traditional DGE and 2) group regularized canonical correlation analysis (GRCCA) – a multivariate regression method that accounts for feature interdependence. Gene set enrichment analyses compared results with established neuropsychiatric risk genes, gene ontology pathways, and cell type enrichments. GRCCA revealed a significant association with SCZ (*P_perm_*=0.001; no significant BD or MDD association), and the resulting gene weight vector correlated with DGE SCZ-control t-statistics (*R*=0.53; *P*<0.05). Both methods indicated down-regulation of immune and microglial genes and upregulation of ion transport and excitatory neuron genes. However, GRCCA - at both the gene and transcript level - showed stronger enrichments (FDR<0.05). Notably, GRCCA results were enriched for SCZ GWAS-implicated genes (FDR<0.05), while DGE results were not. These findings identify a SCZ-specific sgACC gene expression pattern that highlights SCZ risk genes and implicates neuro-immune pathways, thus demonstrating the utility of multivariate approaches to integrate genetic and genomic signals.

## Introduction

Major psychiatric disorders, including schizophrenia (SCZ), bipolar disorder (BD), and major depressive disorder (MDD), are highly heritable^1^, but translating this heritability across genomic levels and into effective treatments remains a critical challenge. Genome-wide association studies (GWAS)^2^ have identified genetic variants robustly associated with psychiatric conditions ^1,3–5^, while case-control differential gene expression (DGE) and weighted gene co-expression network analyses (WGCNA) ^6^ have revealed distinct transcriptomic profiles across these disorders ^7–9^. Despite these advances, gene sets implicated by DGE studies often fail to align with one another or with GWAS findings ^9–11^; leaving the molecular consequences of genetic variation unresolved. This limitation complicates efforts to uncover the molecular underpinnings of psychopathology and develop targeted treatments.

Several factors likely contribute to this lack of agreement between genetic and genomic findings ^12^. Chief among them is the inherent complexity and high dimensionality of genomic data, which encompasses thousands of interrelated features that collectively shape psychopathology. Emerging evidence also highlights the role of alternative RNA splicing, rather than gene expression levels alone, as a key mechanism linking genetic variation to disease ^13–17^. Furthermore, postmortem gene expression is confounded by environmental factors, including medication and recreational drug use ^12,18,19^. Finally, transcriptomic expression patterns are cell-type specific at a granular level and region-specific at a coarser level ^12,17,19–22^, but key studies focus specifically on the prefrontal cortex, particularly the dorsolateral area (dlPFC) ^7,17,23^. These challenges create a complex genomic space that requires thoughtful statistical approaches to derive meaningful insights.

Canonical correlation analysis (CCA) has emerged as a powerful analytic tool for addressing data complexity and high dimensionality in the neuroimaging sphere of biological psychiatry ^24,25^. CCA is a statistical method that identifies associations between two multivariate datasets—such as gene expression and sample covariates—by finding linear combinations of the variables that are maximally correlated ^25,26^. In recent years, CCA has been applied to link brain imaging-derived features in specific brain regions with behavioral measures, psychiatric symptoms, and gene expression patterns ^26^. Group regularized CCA (GRCCA), a recent advancement of CCA, takes into account non-independence of features (for instance, brain regions functioning in networks or gene co-expression) and has been shown to outperform traditional CCA in cross-validation assessments ^27^.

Here, we apply GRCCA to gene expression data derived from post-mortem brain tissue obtained from donors with one of three major psychiatric disorders: SCZ, BD, and MDD. The novel analytical framework we employ aims to extract new insights from existing bulk RNA sequencing data by addressing key limitations of traditional DGE and WGCNA methods. GRCCA distinctly quantifies small, distributed effects across the genome while accounting for interdependence among features (i.e., gene co-expression). As a multivariate approach, it also evaluates the relative contributions of environmental factors—specifically medication and recreational drug use—to observed expression patterns. We apply this method to a deeply sequenced transcriptomic dataset from the subgenual anterior cingulate cortex (sgACC), a limbic region central to mood regulation and underexplored in psychosis research ^8^. The depth of sequencing in this bulk dataset enables us to assess the role of low-expressed genes and alternative transcripts by extending GRCCA to transcript-level analysis. We validate the utility of GRCCA by comparing its results to those from traditional methods (DGE and WGCNA) and evaluating alignment with published risk gene lists. Collectively, our findings demonstrate the utility of multivariate approaches in the transcriptomic space to extract new insights from bulk RNAseq data.

## Methods and Materials

### Samples and RNA sequencing

Analyses included 185 samples (55 Controls, 44 SCZ, 35 BD, and 51 MDD) described by Akula et al^8^ (clinical information is available in **Table S1**). All samples in this study were collected with permission of the next-of-kin under CNS IRB protocols 90M0142 and 17M-N073 or approved by the NIMH Human Brain Collection Core Oversight Committee. Libraries were prepared from total RNA extracted from frozen dissections of sgACC using the RiboZero protocol. Paired end sequencing was performed on Illumina HiSeq 2500. Mapping and quality control were previously described by Akula et al ^8^; reads were mapped to human genome build 38 using Hisat2 and gene and transcript counts were obtained using StringTie ^28^. Here, ‘transcripts’ refers to expressed alternatively spliced gene variants.

### Raw count preprocessing

Genes and transcripts with >= 10 counts across at least 80% of samples were considered in the downstream analysis, resulting in 18,677 genes and 72,403 known transcripts for subsequent analyses. To select informative features and reduce dimensionality ^29^, further filtering was performed on the transcript data. Because mean expression is strongly correlated with variance (*R*=0.97; **Fig S1A**), filtering transcripts based on variance would disproportionately exclude those with low expression levels (i.e., rare transcripts). In contrast, the coefficient of variation (CV) exhibits a much weaker correlation with mean expression (*R*=0.15; **Fig S1B**). Therefore, CV (CV cutoff ∼= 0.36) was used to identify transcripts with minimal variation across samples relative to their mean expression (**Fig S1C**). A total of 54,302 transcripts were included in downstream analyses. After filtering, gene and transcript counts were transformed using variance stabilizing transformation (VST) from DESeq2 package ^30^.

For covariate correction, both known covariates (N=39 total; N=11 technical; N=17 toxicological; **Table S1**) and technical variation due to transcript degradation were considered. Quality surrogate variable analysis (qSVA) was run on VST data to account for transcript degradation ^31^. Although the qSVA transcript degradation matrix was originally derived from the dlPFC, the authors demonstrated the generalizability of the method to other brain regions ^31^. A quality surrogate variable (qSV) was considered significant from the transformed gene counts if it 1) explained greater than 2% of the variance in transcript degradation data (**Fig S2A**) or 2) correlated with known covariates (**Fig S2B**). With these criteria, qSVs 1-7 and 9 were included in the regression. After qSV regression, known covariates that still contributed to variance in the normalized and regressed expression data were identified as those that significantly (FDR < 0.05) correlated with significant (>2% variance explained) gene expression principal components. Age at death and GC percent (a quality control parameter indicating the percentage of RNA bases that are either guanine or cytosine ^32^) met these criteria and were included; sex at birth and race were also included to avoid these variables contributing to module assignments. Together, the following covariates were regressed from the VST counts: qSV1 + qSV2 + qSV3 + qSV4 + qSV5 + qSV6 + qSV7 + qSV9 + sex_at_birth + race + age_at_death + gc_percent. Following these corrections, no significant principal component (PC; > 2% variance explained) was correlated with a technical covariate or significant qSV (**Fig S2C**). The residuals following covariate correction were used for downstream analyses.

### Toxicology multiple correspondence analysis

Seventeen recreational drugs and medications were reported as present, absent, or unknown in the postmortem data based on toxicology reports (**Table S1**; **Fig S3A**). Given the sample size (N=185), the dimensionality of toxicology covariate data was reduced through multiple correspondence analysis (MCA), a method that represents the underlying structure of categorical data ^33^, using the MASS R package (version 7.3-60.2) ^34^. If toxicology data for a given compound was unknown for a given sample, it was assumed not present. The first 8 dimensions of this analysis were selected to represent the toxicology covariate data in subsequent analyses, as each individually explained greater than 5% of variation and collectively they explained the majority of variance (>75%; **Fig S3B**). Compound loadings on each MCA dimension are shown in **Fig S3C**, while known covariate correlations with each MCA dimension can be found in **Fig S3D**.

### DGE analysis

DGE on this dataset was previously run ^8^; however, because raw count preprocessing was updated in the current work, we reran DGE for comparability with WGCNA and GRCCA results. DGE was run on the normalized and corrected expression data using DESeq2 ^30^ version 1.44.0. Toxicology MCA dimensions 1-8 were included as covariates in the model to facilitate direct comparisons with GRCCA results.

### Generation of gene and transcript co-expression modules

The WGCNA R package ^35^ was used to construct gene and transcript co-expression modules. The resulting gene and transcript modules reflect shared underlying biological functionality and/or transcriptional regulation ^36^. In this framework, module 0, termed the ‘gray’ module, corresponds to the set of genes which have not been clustered in any module.

Normalized and corrected expression data for both case and control samples were used to generate co-expression modules. Modules were assigned based on package author recommendations, module size and number, including the gray module, and biological enrichments (**Supplement**; **Fig S4**; **Fig S5**). At the gene level, a soft-thresholding power of 3, a minimum module size of 40, and a tree cut height of 0.980 were used, resulting in 23 co-expression modules (**Fig S4**, **Fig S5A**). For the transcript expression data, a soft-thresholding power of 2, a minimum module size of 35, and a tree cut height of 0.988 were used, resulting in 40 transcript-level modules.

### Characterizing the enrichment of co-expression modules

Biological relevance of gene and transcript modules was assessed using 2 distinct enrichment tests:

<u>Functional enrichment</u> of each of the modules (including biological process (BP), cellular component (CC), and molecular function (MF)) was assessed using the enrichGO function from the R package clusterProfiler ^37^. To perform this analysis on transcript-level results, transcripts were mapped to genes using the biomaRt package in R version 2.60.1 ^38^. To summarize the functional enrichment results, topic modeling was employed on all significant gene ontology (GO) terms across all modules ^39^(**Supplement**; **Fig S5B**).

<u>Cell-type enrichment</u> of each module was calculated as the hypergeometric overlap between the genes in a module and genes associated with each cell type, as annotated by Lake et al ^40^ and grouped into seven broader cell-type classes by Seidlitz et al ^41^ (**Fig S5C**). Transcripts were mapped to genes to perform this analysis at the transcript level.

### Developmental trajectory analysis

To assess the co-expression patterns of genes through development, we determined the average developmental trajectory for each module. Neocortical gene expression values across different windows of life from 421 samples from 41 human brains were accessed from PsychENCODE data ^42^. For each module, genes were averaged across samples at each time window and plotted as a smooth curve to visualize periods of average higher and lower expression (**Fig S5D**). Transcripts were mapped to gene level to perform this analysis at the transcript level.

### Canonical correlation analysis (CCA)

CCA is a technique that determines the linear association between two multivariate data matrices from different modalities ^25^. Here, we used a custom version of the CCA/PLS toolkit [https://github.com/rlsmith1/sgACC_transcriptomics_analyses/tree/main/RCCA_toolkit/cca_pls_toolkit_final] ^26^, with the *X* matrix containing samples-by-expression data (N = 18677 genes; N = 54302 transcripts), and the *Y* matrix samples-by-covariates, including psychiatric diagnosis and 8 toxicology latent dimensions (**Fig S4**).

The output of CCA is a vector of feature (gene/transcript) weights (*w_x_*) and a vector of covariate weights (*w_y_*), which are the coefficients used to construct the X and Y latent variables (*LV_x_*=*X*⋅*w_x_* and *LV_y_*=*Y*⋅*w_y_*, respectively) (**Fig S6**; **Table 1**). These weights are optimized by the CCA algorithm to maximize the correlation between the latent variables. To assess the consistency of these weights across 1000 bootstraps, a *Z*-score was calculated for each gene/transcript and covariate weight as the actual weight divided by its standard deviation. To estimate feature contribution to the identified association, we used structure correlations, defined as the Pearson correlation between each feature (i.e., matrix column) and its corresponding latent variable (cor(*X*, *LVx*) and cor(*Y*, *LVy*), respectively; **Table 1**) ^26^. In this work, dual criteria were applied to determine feature significance: (i) |*Z*| >= 2, to ensure the weight remained consistently non-zero across bootstraps, and (ii) structure correlation (*r*_x_) FDR < 0.05, to confirm the variable’s correlation with the *X*-*Y* associative effect.

**Table 1.** (G)RCCA model inputs and outputs.

|  |  |  |  |
| --- | --- | --- | --- |
| **Inputs** | **Data** | $X$ | $Y$ |
|  | **\*Variance explained** | 0.1:1:0.1 |  |
|  | **†Group vector** | WGCNA module | - |
|  | **†Group hyperparameter (mu)** | 0.1 | - |
|  | **Feature hyperparameter (lambda)** | 1-1/n features | - |
| **Outputs** | **Weight** | $w_x$ | $w_y$ |
| | **Latent variable** | $LV_x = X \cdot w_x$ | $LV_y = Y \cdot w_y$ |
| | **Structure correlation** | $r_x = \text{cor}(X, LV_x)$ | $r_y = \text{cor}(Y, LV_y)$ |

\*Can be value or search space
†GRCCA only

### Regularized CCA (RCCA)

In analyses where the sample size is smaller than the number of variables, a standard CCA model is ill-posed (i.e., it doesn’t have a unique solution). Regularization (i.e., treating all canonical coefficients equally and shrinking them to zero) and dimensionality reduction successfully address this issue ^26^. Thus, we (1) included a regularization parameter for the *X*-matrix defined by the number of features (lambda = 1-1/(N features)), and (2) optimized the amount of variance in the *X* matrix that was incorporated in the model (search space from 0.1 to 1 by increments of 0.1) using a permutation-based approach to avoid overfitting^43^.

### Group RCCA (GRCCA)

A limitation of the standard RCCA approach is that it ignores underlying data structure and treats all features equally ^27^. However, in the case of transcriptomics, this is an incorrect assumption as genes and transcripts are co-expressed. Thus, it is further useful to regularize at the group level, in addition to the feature level. Here, we use WGCNA module assignment as the grouping vector. The group-level regularization parameter was set to mu = 0.1. The feature-level regularization parameter and the variance explained search space were consistent between RCCA and GRCCA algorithms to maximize comparability (refer to the **Supplement** for RCCA results). See **Table 1** for (G)RCCA inputs and outputs.

### Gene set validation and characterization (gene set enrichment analyses; GSEA)

To test the biological validity of GRCCA and compare it with DGE, analysis results were benchmarked using published gene lists and functional enrichment tests. GSEA was run using the R package fgsea (version 1.30.0)^44^ to determine the rank-based enrichments of analysis result distributions of the following: (1) neuropsychiatric risk genes (SCZ, BD, MDD, and Autism Spectrum Disorder (ASD); **Supplement**), (2) cell type ^40^, and (3) GO functional pathways (including BP, CC, and MF ontologies). Per GSEA author recommendations, the full vectors of gene DGE t-statistics and structure correlations were used as input (i.e., no filtering was applied).

In all statistical and enrichment analyses, *P* values were corrected for multiple comparisons using the Benjamini-Hochberg (BH) method.

Code for all analyses is available at https://github.com/rlsmith1/sgACC_transcriptomics_analyses.git. An overview of the study analysis pipeline is available in **Figure 1**.

**Figure 1.**
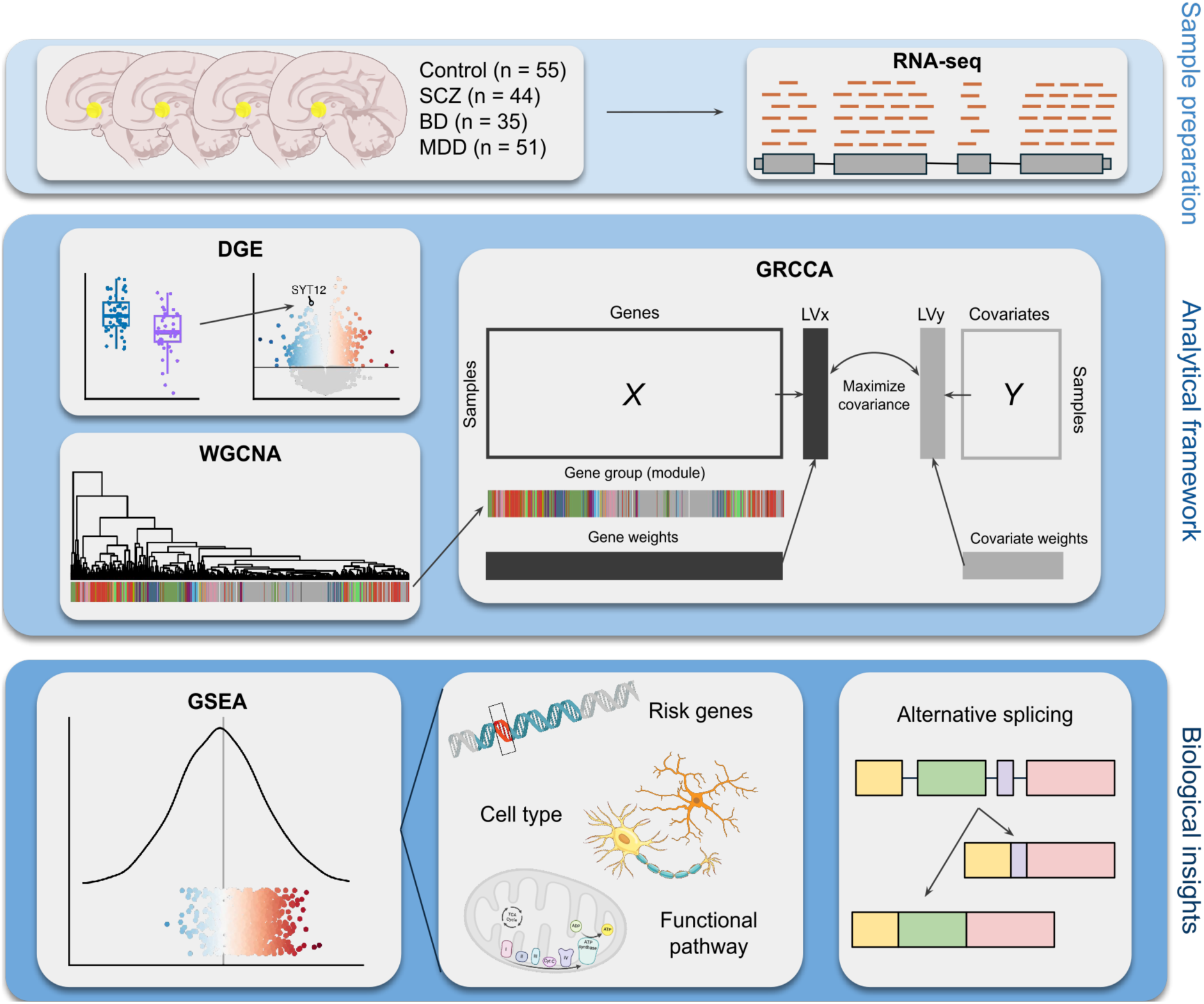
A schematic overview of the analysis pipeline. Abbreviations: SCZ = schizophrenia; BD = bipolar disorder; MDD = major depressive disorder; DGE = Differential gene expression (analysis); WGCNA = weighted gene co-expression network analysis; GRCCA = group regularized canonical correlation analysis; GSEA = gene set enrichment analysis.

## Results

***GRCCA identified a robust link between gene expression and schizophrenia*** The GRCCA model was optimized by incorporating 70% of gene expression variance, resulting in a significant X-Y latent variable correlation of 0.585 with *P*=0.001 across 1000 permutations (**Fig 2A**; **Table S2A**). One latent variable was significant and is reported here. Schizophrenia was the only covariate significantly associated with this latent variable, with a *Z*-score of 3.50 (**Fig S7**) and a structure correlation *r*_y_=0.921 (FDR<0.001; **Fig 2B**; **Table S2B**; see **Supplement** and **Fig S6** for an explanation of CCA terminology and design). Thus, the covariate association with gene expression in this latent variable was primarily driven by SCZ.

**Figure 2.**
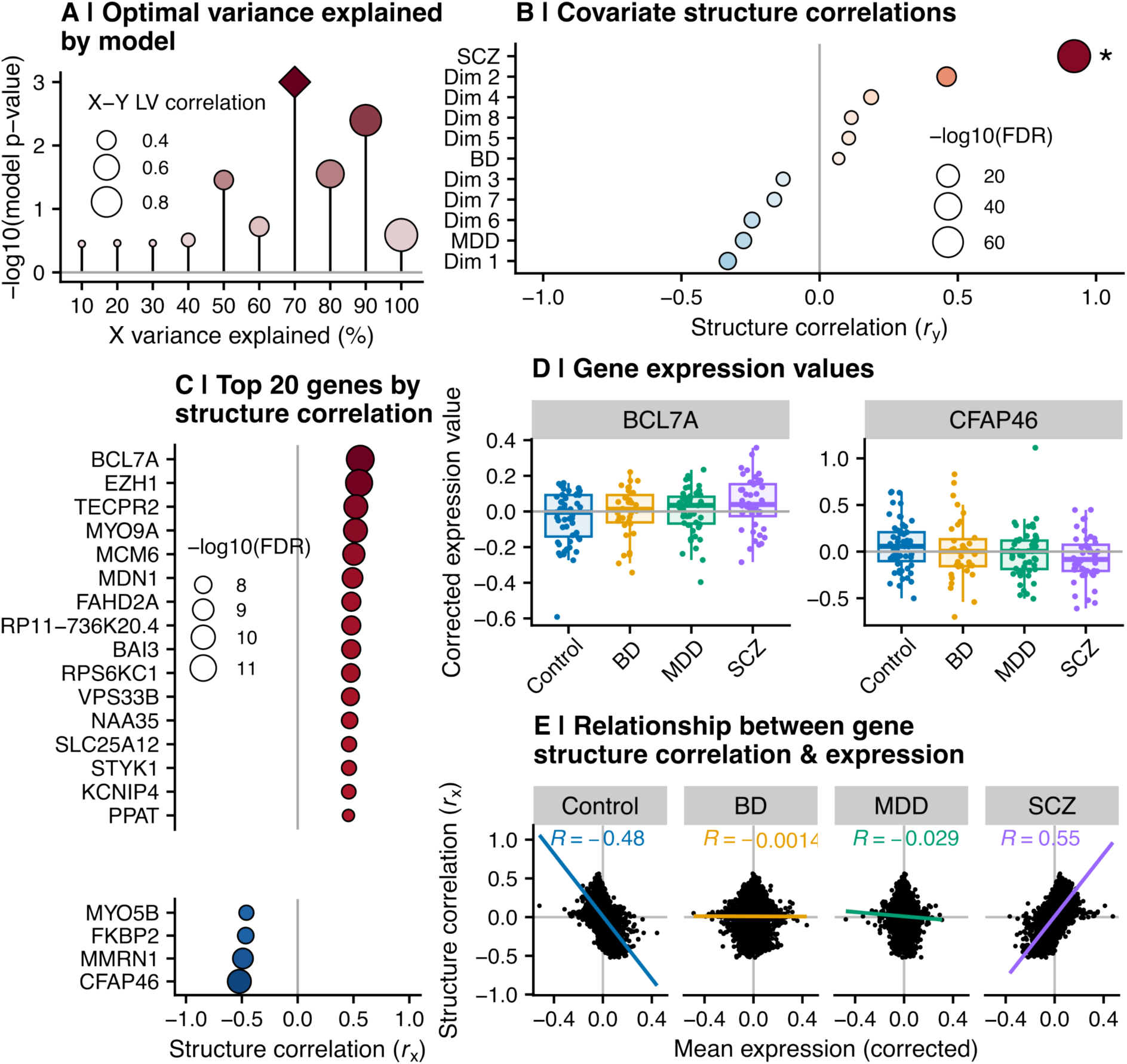
GRCCA identified a linear association between SCZ and gene expression. **(A)** GRCCA model *P* values by percent variance explained in the X matrix (gene expression data). Point size indicates X-Y latent variable correlation, while the color indicates -log10(*P*-value). As *P* values were calculated across 1000 bootstraps, the minimum *P* value possible is 0.001 (-log10(*P*) = 3; indicated by diamond). **(B)** Covariate structure correlations, calculated as the Pearson correlation between covariate value across samples and the Y latent variable (U). The x-axis indicates structure correlation (*r*_y_), while the y-axis represents each covariate, ordered by increasing structure correlation. Point color also indicates structure correlation, while point size represents correlation significance -log10(FDR). **(C)** The top 20 genes subset by the absolute value of their structure correlation (*r*_x_). Each gene is represented on the y-axis, and their respective structure correlations are indicated by the x-axis and point color. Point size represents correlation significance - log10(FDR). **(D)** Expression values across diagnostic groups for the genes with the highest (BCL7A, left) and lowest (CFAP46, right) structure correlations. Each point is a sample, colored and ordered on the x-axis by diagnostic group. The y-axis shows the normalized & correlated expression value for that sample. The box and whiskers show the distribution of values for each diagnostic group. Neither gene was differentially expressed per the current study’s DGE analysis (*p* BCL7A = 0.118; *t* BCL7A = 1.56; *p* CFAP46 = 0.065; *t* CFAP46 = -1.85). **(E)** The correlation between mean gene expression and structure correlation, faceted by diagnostic group. Each point is a gene; its position on the x-axis indicates the mean expression value across samples within the diagnostic group, while the y-axis represents its structure correlation (*r*_x_). The line shows the line of best fit, colored by diagnostic group, and inset text indicates the correlation within each diagnostic group.

According to the same criteria (FDR<0.05 and |*Z*|>=2), 1,211 genes were significantly associated with the latent variable correlated with SCZ (**Fig S8**; **Table S2C**). The top 20 by |*r*_x_| are shown in **Fig 2C**. The gene with the highest structure correlation, BCL7A (*r*_x_=0.56; also a SCZ risk gene), showed higher expression in SCZ compared to controls (**Fig 2D, left**), but was not significantly differentially expressed according to standard DGE analysis (*P*=0.12; L2FC=0.01). Similarly, the gene with the lowest structure correlation, CFAP46 (*r*_x_=-0.52), demonstrated lower expression in SCZ compared to controls (**Fig 2D, right**), but was not differentially expressed (*P*=0.06; L2FC=-0.02). This pattern holds across genes: genes with higher expression levels in SCZ have positive *r*_x_ and vice versa for lower expression and negative *r*_x_ (*R*=0.55; *P*<0.001; **Fig 2E**); while the reverse is true for controls (higher expression levels are associated with lower or more negative *r*_x_; *R*=-0.48; *P*<0.001). There are no significant relationships between gene expression in BD and MDD and structure correlation, providing further evidence that the association we detected was driven by SCZ.

### SCZ risk genes were specifically and significantly enriched in the GRCCA gene structure correlation distribution

To test the alignment of GRCCA results with genetic variants, we tested the structure correlation distribution for overrepresentation of known SCZ risk genes based on common or rare variant association studies ^3,45^. To assess specificity to SCZ, we also ran the same enrichment analysis using known risk genes for autism spectrum disorder (ASD) ^46^, MDD ^5^, and BD ^4^ (all based on common variant associations). The positive end of the structure correlation distribution was significantly enriched for SCZ common-variant associated genes (both the broad set normalized enrichment score, NES=1.57; FDR=1.1*10^-4^) and the prioritized gene list (NES=1.54; FDR=6.5*10^-3^); **Fig 3A**; **Table S3A**). SCZ rare variant-associated genes were also positively, though not significantly, enriched within the structure correlation distribution (NES = 1.36; FDR = 0.18). In contrast, risk genes associated with ASD, BD, and MDD were not enriched in either end of the structure correlation distribution (ASD: FDR=0.45, NES=1.00; BD: FDR=0.37, NES=1.08; MDD: FDR=0.10, NES=1.26). Taken together, these results suggest that genes with positive structure correlations are significantly enriched for SCZ risk genes, and that this signal is specific to SCZ and not other psychiatric disorders.

**Figure 3.**
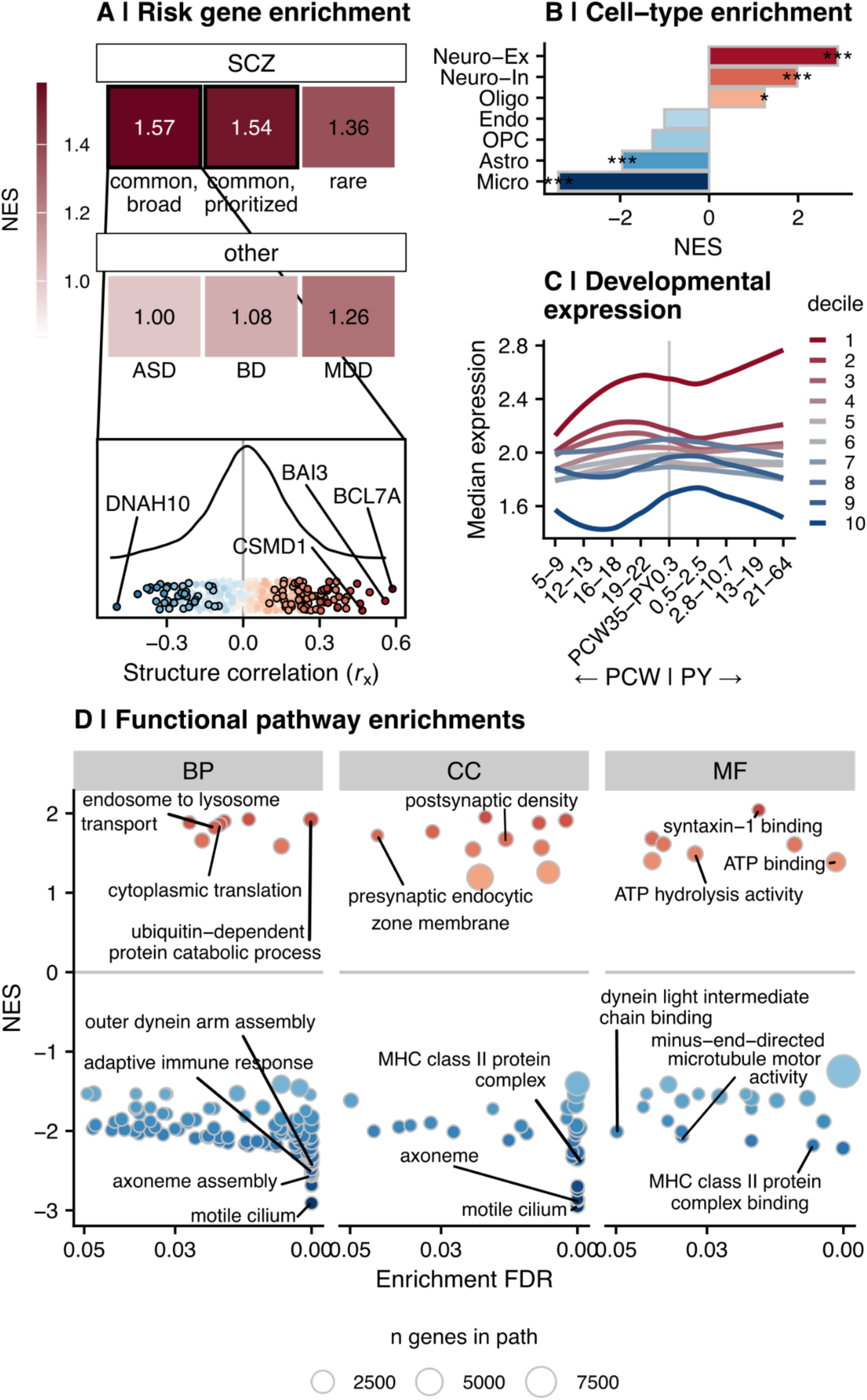
GRCCA identified a neuro-immune gradient of gene expression associated with SCZ. **(A)** Enrichment of SCZ-control DEGs for risk genes identified in psychiatry GWAS. **Top:** Tile color and text shows the normalized enrichment score (NES) for each association; significant enrichments (FDR < 0.05) are outlined in black. The top row shows SCZ risk genes (broad fine-mapped common variant-associated genes ^3^ (N = 628; FDR = 1.1*10^-4^); prioritized common variants ^3^ (N = 120; FDR = 6.5*10^-3^;); rare variants ^45^ (N = 10; FDR = 0.18)), while the bottom row shows risk genes identified through GWAS of other psychiatric disorders: (Autism Spectrum Disorder (ASD) common variants ^46^ (N = 567; FDR = 0.45); BD common variants ^4^ (N = 162; FDR = 0.37); MDD common variants ^5^ (N = 339; FDR = 0.10)). **Bottom:** Position of SCZ common variant-associated risk genes (broad set) in the GRCCA structure correlation distribution. The curve shows the density distribution of the structure correlation (*r*_x_) of DEGs in the current study, while the points show the position of SCZ risk genes in the distribution. Points are colored by structure correlation, where red indicates positive correlation with SCZ and blue indicates negative correlation with SCZ. Significant GRCCA genes (|*Z*| >= 2; *r*_x_ FDR < 0.05) are outlined in black, while top significant risk genes by |*r*_x_ | are labeled. **(B)** Cell type enrichment of GRCCA results by gene set enrichment analysis (GSEA). The y-axis shows each cell type, while the x-axis and bar color indicate the GSEA normalized enrichment score (NES). A negative NES (blue) indicates that genes associated with the cell type (per ^40^) are enriched at the negative end of the structure correlation (*r*_x_) distribution, while a positive NES indicates the same for the positive end. Significance is indcated by asterisks as follows: * FDR < 0.05; ** FDR < 0.01; *** FDR < 0.001. **(C)** Developmental trajectories of genes in each GRCCA decile per the PsychENCODE data ^42^. The x-axis shows the developmental window, split by post-conception week (PCW) and post-natal year (PY). The y-axis shows the median PsychENCODE expression value for samples from the frontal cortex. Each line represents a GRCCA decile (binned according to structure correlation (*r*_x_)); with red representing the top decile (i.e., most positive structure correlations) and blue representing the bottom decile (i.e., most negative structure correlations). **(D)** Gene ontology gene set enrichment analysis (GSEA) results for the GRCCA structure correlation (*r*_x_) distribution. Genes were ranked by *r*_x_, and GSEA determined GO pathway enrichments at either end of the distribution (positive or negative). Each facet represents a distinct GO ontology (BP = biological process, CC = cellular component, MF = molecular function). The x-axis shows the -log_10_(FDR) of the pathway, while the y-axis indicates its NES, in which a positive value indicates the pathway was significant in genes with a positive structure correlation, while a negative value indicates the pathway was significant in genes with a negative structure correlation. Points are colored by NES and sized by the number of genes in the pathway.

### The distribution of gene structure correlations represented a developmentally sensitive, polarized axis of neuron-immune enrichments

We then assessed the biological enrichments of the vector of gene structure correlations (*r*_x_) using GSEA. Cell-type GSEA revealed an overrepresentation of genes expressed in excitatory neurons (NES=2.89; FDR<0.001) and inhibitory neurons (NES=1.98; FDR<0.001) at the positive end of the structure correlation distribution (i.e., genes that tended to be upregulated in SCZ) (**Fig 3B**; **Table S3B**). In contrast, the negative end of the distribution was enriched for genes expressed in microglia (NES=-3.39; FDR<0.001) and astrocytes (NES=-1.95; FDR<0.001), revealing an enrichment pattern with neurons at one pole and glia at the other pole. Functional pathway GSEA aligned with these cell type results: The positive end of the structure correlation distribution contained genes associated with synaptic signaling, ubiquitination, and vesicular transport, typical of neurons. Among genes with negative structure correlations, pathways corresponding to immune response and cilium movement/assembly–typical glial functions–were strongly enriched (**Fig 3D**; **Table S3C**). These biological enrichments suggest that the GRCCA structure correlation distribution represents a synaptic-immune gradient of expression that is differentially regulated in SCZ.

Based on convergent evidence that SCZ is a neurodevelopmental disorder ^47^, we hypothesized that the genes identified through this analysis are developmentally sensitive. To test this, we split the vector of structure correlations into deciles, with genes at the positive end in decile 1 and genes at the negative end in decile 10. We then summarized the normative developmental curves of each decile using PsychENCODE neocortical data ^42^. Genes in the first decile had high levels of expression that increased through development, with a local maximum right before birth (**Fig 3C**). Genes in the tenth decile had low levels of expression and peak expression immediately after birth. In contrast, genes in deciles 2-9 showed less striking levels or patterns of expression through development. Together, these results suggest that genes that are most strongly associated with SCZ (either positively or negatively) show dynamic developmental patterns of expression, especially around time of birth.

### Differential gene expression (DGE) analysis results demonstrated weaker biological enrichments and did not align with SCZ risk genes

To compare our novel use of GRCCA with current state-of-the-art methods, we ran standard differential gene expression analysis on the preprocessed expression data. 1,690 genes were identified as differentially expressed between SCZ and controls at *P*<0.05 (DEGs; **Fig 4A**; **Fig 4B**; **Table S4**; five DEGs at FDR<0.05). Of these, 433 were also identified as DEGs (*P*<0.05) in a previous differential expression analysis of these data (total N DEGs=1373) ^8^ (**Fig S9A**; hypergeometric *P* < 0.001; odds ratio = 2.30), indicating statistical preservation across studies with distinct preprocessing pipelines. Furthermore, the differential expression *t*-statistic across genes was significantly correlated with the *t*-statistics reported in Akula et al ^8^ (**Fig S9B**; *R*=0.61; *P*<0.001). DGE effect size was also correlated with GRCCA gene structure correlation (*r*_x_; *R*=0.53; *P*<0.001; **Fig 4C**; **Fig S9A**), demonstrating that GRCCA structure correlation is indicative, but independent, of up- or down-regulation per DGE analysis.

**Figure 4.**
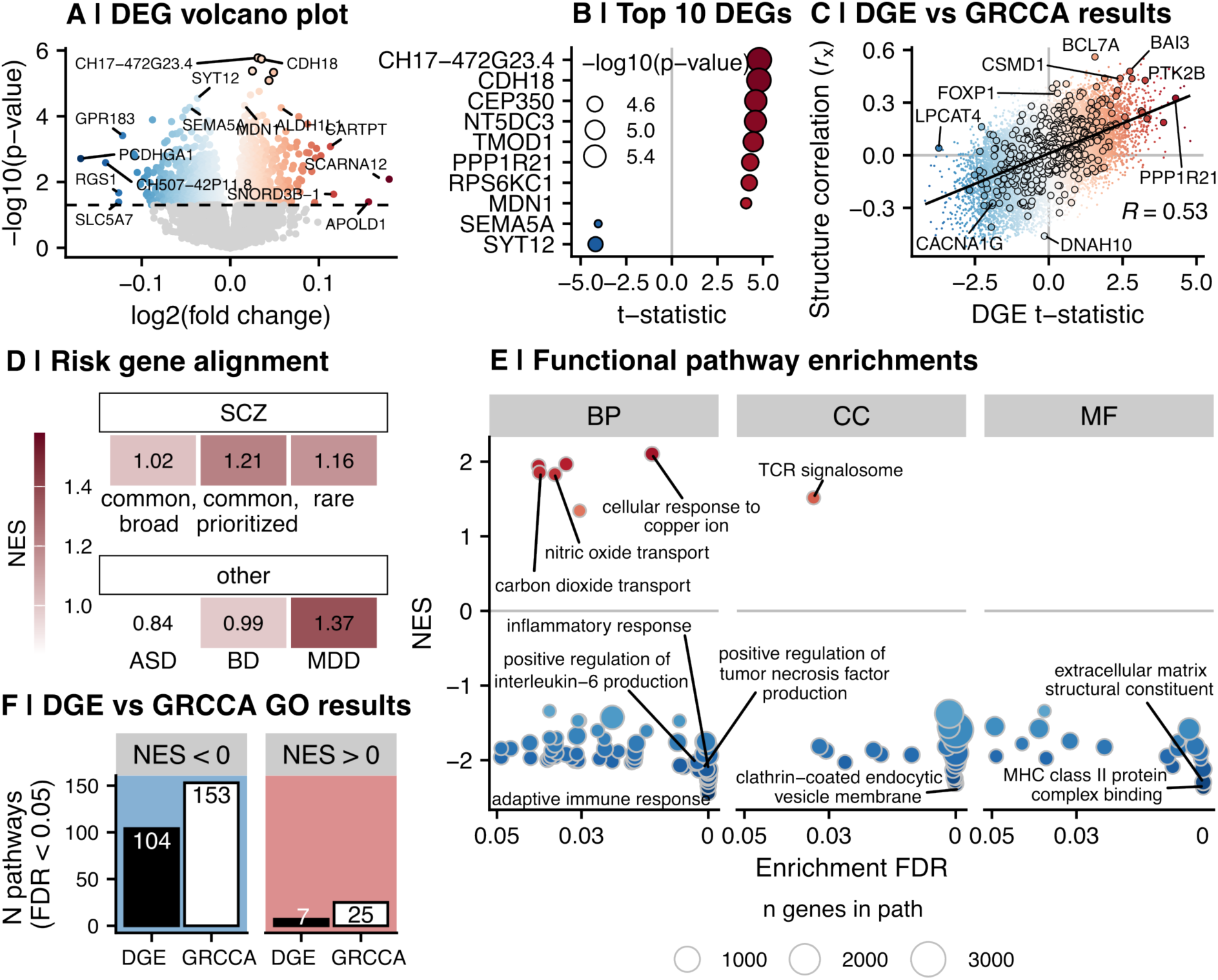
A comparison of traditional SCZ-control DGE analysis and GRCCA results. **(A)** Volcano plot showing SCZ-control differential gene expression in the current analysis. The x-axis represents the log2(fold change) (L2FC) of the gene, in which a positive change (red) indicates the gene was upregulated in SCZ, and a negative change (blue) indicates the gene was downregulated in SCZ. The y-axis shows the -log10(p-value), and genes that were significant after FDR correction are outlined in black. The dashed line indicates *P*=0.05; genes that did not meet this threshold are colored in gray. Genes with low p-values (-log10(*P*) > 4.2) and/or high absolute fold change (abs(L2FC) > 0.11) are labeled. **(B)** The 10 differentially expressed genes (DEGs) with the highest absolute value effect size (*t*-statistic) in the DGE analysis. The y-axis lists the symbols for these genes, while the x-axis & point color indicate their respective effect sizes. Point size indicates - log10(*P*). **(C)** Relationship between gene structure correlation (*r*_x_) and DE *t*-statistic from the differential gene expression analysis. The x-axis shows the DE effect size for each gene and the y-axis shows their respective structure correlations, determined by GRCCA. Points are colored by structure correlation; risk genes are outlined in black. The DE *t*-statistic and GRCCA *r_x_* are correlated at *r* = 0.53 (*P* < 0.001). **(D)** Enrichment of SCZ-control DEGs for risk genes identified in psychiatric disorder-related GWAS. Panel legend is the same as Fig 3A, with the following statistics: SCZ common (broad) *P* = 0.48; SCZ common (prioritized) *P* = 0.33; SCZ rare *P* = 0.48; ASD *P* = 0.96; BD *P* = 0.56; MDD *P* = 0.07. **(E)** Gene ontology gene set enrichment analysis (GSEA) results for the distribution of DGE effect sizes. Panel legend is the same as Fig 3D. **(F)** The number of significant GO pathways identified by DGE and GRCCA. The x-axis and bar color indicates analysis (DGE or GRCCA), and the height of the bar on the y-axis shows the number of significant pathways per GO GSEA (FDR < 0.05). The plots are facetted by pathway direction, where negatively enriched pathways (NES < 0.05; blue) is represented on the left and positively enriched pathways (NES > 0.05; red) is represented on the right.

DGE analysis resulted in a vector of case-control *t*-statistics (in which a positive *t*-statistic indicates upregulation in SCZ) across genes. To assess DGE alignment with genetic variation identified in psychiatric disorders, we tested the SCZ-control *t*-statistic distribution for SCZ, ASD, MDD, and BD risk gene list enrichment (**Fig 4D**; **Table S5C**). Though all risk gene lists tended to cluster at the positive end of the effect size distribution, none of the enrichments were significant (all *P*>0.05). The strongest enrichment was for MDD, which showed a positively skewed but non-significant clustering of risk genes in the SCZ-control effect size distribution (NES=1.37; FDR=0.07; **Fig 4F**; **Table S5C**). Thus, unlike GRCCA, DGE failed to align with SCZ genetic variation.

Finally, to directly compare biological information contained across analysis results, we ran the same enrichment tests on this DGE vector as with the GRCCA structure correlations. Functional pathway GSEA demonstrated that, as with GRCCA, the negative end of the distribution (i.e., genes downregulated in SCZ compared to controls) was strongly enriched for immune processes (**Fig 4E**; **Table S5B**). The positive end of the effect size distribution was only weakly enriched for genes in molecular and ion transport pathways. For both the negative and positive end of the gene results distributions, GSEA identified more significant enrichments (at FDR<0.05) in the GRCCA results than in the DGE results (**Fig 4F**), demonstrating stronger biological enrichment of GRCCA results.

#### Transcript-level analysis revealed isoform-specific patterns for SCZ risk genes

Increasing evidence supports dysregulation in alternative splicing patterns as a key link between genetic variation and neuropsychiatric disease ^14–16,21^. Thus, we leveraged this deeply sequenced dataset and ran the same GRCCA model on the normalized and corrected transcript-level expression data (N=54302; **Fig S10A**). SCZ once again emerged as the covariate most strongly associated with the latent variable (*r_y_*=0.98; FDR<0.001; **Fig S10B**); however, the overall model did not reach significance at *P*=0.001, likely due to the high number of features relative to sample size (*P*=0.009; **Fig S10A**). Therefore, to better illustrate the known SCZ signal, we re-ran the model including only transcript derivatives of either: (a) genes associated with common variants linked to SCZ, BD, MDD, or ASD, or (b) genes significantly associated with the latent variable in the gene-level GRCCA (*r_x_* FDR<0.05) (collective N=12986). In this subset analysis, the best fitting model incorporated 40% of the variance in the expression data, optimizing the X-Y latent variable correlation at 0.67 and *P*=0.001 across 1000 permutations (**Table S6A**; **Fig S10C**). SCZ maintained its salient latent variable association (*r_y_*=0.94; FDR<0.001; **Fig 5A**, **Table S6B**). Covariate and transcript structure correlations are provided as a resource in this paper (**Table S6B** and **Table S6C**).

**Figure 5.**
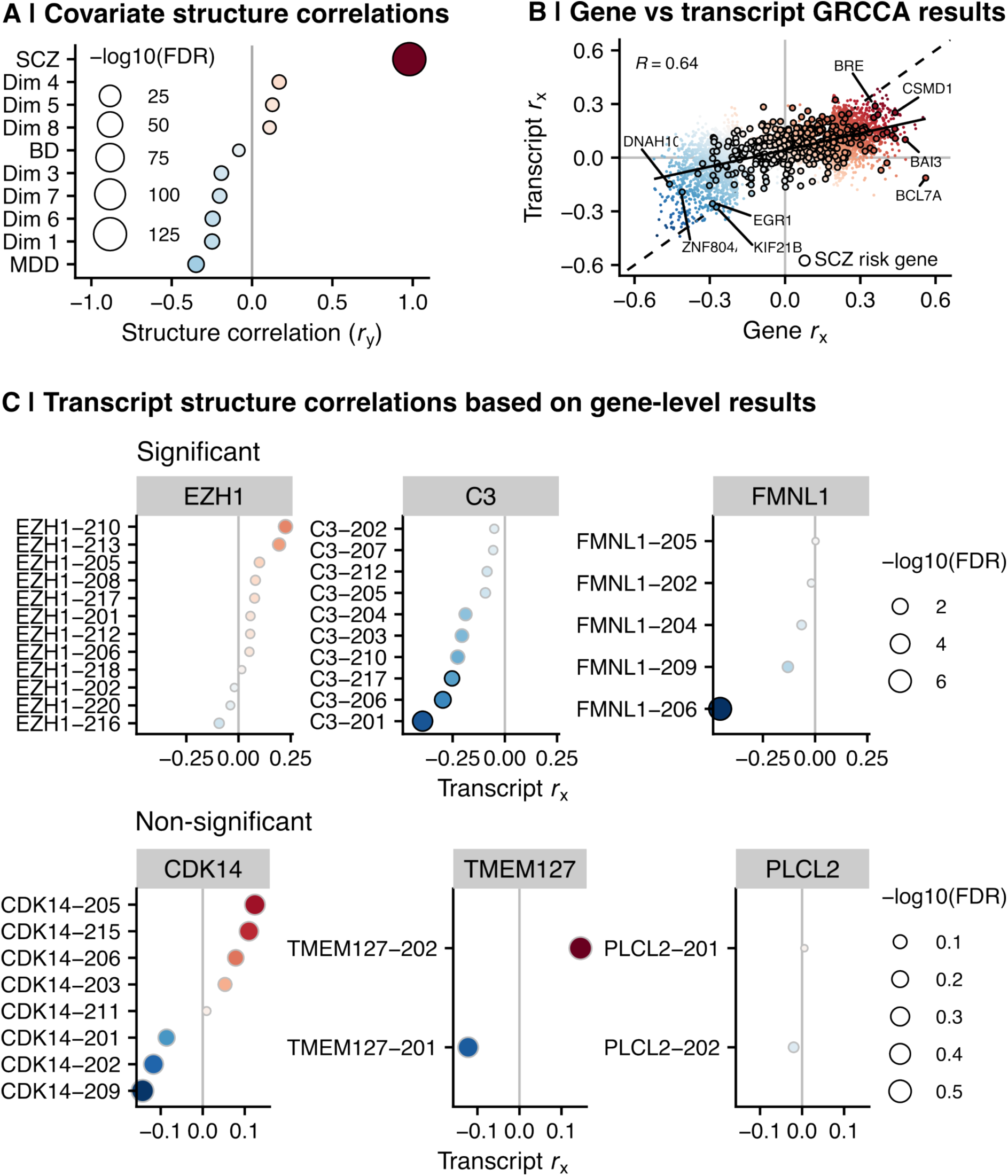
Alternative splicing patterns revealed by transcript-level GRCCA. **(A)** Covariate structure correlations (*r*_y_) as determined by transcript-level GRCCA. Panel legend is the same as **Fig 2B**. **(B)** Gene-level structure correlations (x-axis) and transcript-level structure correlations (y-axis) were correlated at *r* = 0.64 (*P* < 0.001). Transcript structure correlations were mapped to gene-level by taking the maximum absolute structure correlation across all transcript derivatives of a specific gene. SCZ risk genes are outlined in black. **(C)** Structure correlations of transcript derivatives of select SCZ-related genes that were significant (top) and non-significant (bottom) in the gene-level GRCCA analysis.

Transcript structure correlations were significantly correlated with gene structure correlations (*R*=0.64; *P*<0.001; **Fig 5B**), demonstrating alignment across analysis levels. In general, SCZ risk genes that were positive at the gene-level were also positive at the transcript level (**Fig 5B**). However, transcript derivatives of the same gene demonstrated distinct structure correlation patterns, giving rise to gene-level significance (or non-significance). A gene could be identified as significant if it had one or more significant transcripts driving the effect (e.g., in the case of FMNL1 and C3; **Fig 5C**), or an accumulative effect of several sub-threshold transcripts in the same direction (e.g., EZH1). In contrast, the transcript derivatives of many non-significant genes were also not significant (e.g., PLCL2; **Fig 5C**); however, some non-significant genes did have one or more significant transcript derivatives with opposite directions of effect (e.g., TMEM127 and CDK14). These findings underscore the importance of analyzing transcript-level data to identify expression patterns that cannot be detected at the gene-level.

## Discussion

Complex psychiatric disorders are marked by subtle, coordinated changes across many genes, making such disorders particularly well-suited to network-based multivariate analyses. Here, we adapted a multivariate approach, GRCCA, previously used in neuroimaging, to characterize the transcriptional landscape of sgACC in three major mental disorders. The results capture gene expression patterns often missed by traditional methods.

Currently, there are two widely used univariate approaches to characterize gene expression patterns in post-mortem brain tissue. Differential gene expression analysis identifies individual genes with statistically different levels of expression across conditions ^48^. This approach fails to capture the biological interdependencies among genes, which are known to function in coordinated programs. WGCNA accounts for this limitation by constructing gene networks based on expression patterns across samples^6^ and defining modules that reflect shared underlying biological functionality and/or transcriptional regulation^36^. However, WGCNA comes with its own limitations, as functionally fluid gene modules are categorically assigned. Also, current methods linking modules to covariates of interest (e.g., psychiatric disorder) ignore a large proportion of variance in expression data, as only the first principal component (PC1) of the module expression matrix is considered ^35,49^. Though useful, DGE and WGCNA have thus far not implicated a convergent gene set across schizophrenia studies, nor do they consistently align with gene sets that have been curated by GWAS ^9^.

GRCCA, a multivariate approach, addresses the limitations of both traditional approaches by modeling distributed effects across genes and covariates. It additionally accounts for nonindependence of features by leveraging information about the underlying data structure (in this case, WGCNA-derived co-expression modules). Unlike WGCNA alone, GRCCA incorporates optimal variance in the expression data (i.e., more than PC1 without overfitting). Altogether, GRCCA accounts for interactions of genes with each other and with environmental covariates, including medications.

In this GRCCA analysis, one significant latent, or “hidden”, variable was identified, representing the shared variation between gene expression and all covariates (including both psychiatric disorders and toxicology data). The covariate with by far the most significant structure correlation, and thus highest contribution to the latent variable, was SCZ. This contribution was above that of MDD or BD, or any of the toxicology dimension covariates. Detangling the effects of the many factors that drive gene expression, notably medication and recreational drug usage, is an outstanding challenge in human post-mortem transcriptomic studies ^9,11,12,18^. Though multivariate GRCCA does not measure the causality of disorder-related changes in gene expression versus consequential and environmental factors accumulated over the lifetime, it is able to quantify relative contributions of these variables, and thus is a step towards decoupling their respective effects.

The significant and specific enrichment of SCZ risk genes in the GRCCA results provides strong evidence of such disorder-covariate decoupling. Common variant-associated genes identified through SCZ GWAS ^3^ were overrepresented at the positive end of the GRCCA gene structure correlation vector. In contrast, genes associated with ASD, BD, and MDD were not; a finding that is potentially influenced by the lower SNP-based heritability estimates of these phenotypes compared to SCZ ^3–5,46^. It is unsurprising that SCZ risk genes were clustered toward the positive end of the GRCCA results, as this pole was enriched for neurons and synaptic signaling pathways, and GWAS association signals with SCZ are known to be highly enriched in neurons ^1,3^. To our knowledge, this risk gene enrichment is a unique finding in transcriptomic analysis of bulk post-mortem tissue ^9,11^. Conversely, it was unexpected to find rare variants nominally enriched among genes with positive structure correlations; however, previous studies have shown that the downstream expression impacts of loss-of-function genetic mutations are complex and varied, highlighting the role of compensatory mechanisms ^50,51^. It is of further note that SCZ-related variation in gene expression, even once decoupled from environmental covariates, may represent a lifetime consequence rather than an underlying cause of the disorder^12^. However, in demonstrating that genes with allelic variants associated with SCZ also show robust expression changes in our GRCCA analysis, we highlight a clear connection across biological levels in SCZ pathophysiology.

Another strength of the GRCCA approach is its biological interpretation, as the GO pathway enrichments of the gene structure correlation vector were stronger than the pathway enrichments of the gene *t*-statistic vector determined via DGE analysis. GSEA of GO terms revealed an upregulation of genes associated with neurons and neuron-related pathways, including vesicle transport and synaptic signaling, and downregulation of genes associated with glial cells and immune-related and cellular transport pathways. Post-mortem analyses of the schizophrenia transcriptome currently provide mixed evidence for the role and direction of neuron- and immune-related enrichments ^52^. For example, a recent SCZ single-cell study provides evidence for downregulation of synaptic signaling genes in excitatory neurons ^53^. In transdiagnostic bulk data, Gandal and colleagues described a broad gradient of dysregulation with the down-regulation of neuronal and synaptic signaling genes and the up-regulation of glial-immune signals ^7^. However, specific to SCZ, they indeed report down-regulation of a microglia-specific gene module, and up-regulation of neuron and synaptic signaling/vesicle modules ^7^. A cross-regional study identified multiple immune gene sets that were down-regulated in the dlPFC and hippocampus in SCZ ^54^. A previous study in the ACC also identified down-regulation of immune-associated genes in SCZ, and the upregulation of ubiquitin-proteasome system genes (congruent with our results) ^55^. Our findings thus contribute to a growing literature implicating the dysregulation of immune- and neuron-related genes in SCZ.

A key limitation in the interpretation of the GRCCA results is that they were derived from bulk, rather than single-cell data. Cell-type correction was not performed as a preprocessing step due to lack of a well-validated deconvolution method ^56^. Thus, the apparent decrease of immune-related genes in SCZ could reflect a decrease in the proportion of (micro)glia, rather than down-regulation of the genes themselves. However, available single-cell data do not support a substantial change in any major cell-type fraction in SCZ ^53^; thus, it is plausible that the enrichments we detected actually reflect a change in gene expression. Furthermore, the GRCCA vector of gene structure correlations was significantly anticorrelated with a single-cell-derived gene expression latent factor shown to decrease in SCZ (**Fig S11**) ^57^, demonstrating consistency of bulk-derived GRCCA results with single-cell findings.

In sum, we illustrate a promising new approach to characterizing disorder-related gene expression differences in post-mortem transcriptomic data. Due to the inherent complexity of genomic data, methods that can adequately handle such complexity are required to characterize the molecular landscape of psychiatric disorders. We have presented evidence for a developmentally-sensitive, neuro-immune gradient of transcriptomic dysregulation in schizophrenia. We have published the gene- and transcript-level datasets and computational tools used to generate these results as an open resource to facilitate usage across post-mortem transcriptomic studies.

## Data Availability

The raw count data can be downloaded from dbGAP at https://www.ncbi.nlm.nih.gov/projects/gap/cgi-bin/study.cgi?study_id=phs000979.v2.p2. The case-control differential gene expression results reported by Akula et al (2021) ^8^ can be found at https://www.nature.com/articles/s41386-020-00949-5 (Supplementary Table 4 for SCZ vs controls). PsychENCODE developmental expression data are made available by Li et al (2018) ^42^ https://www.science.org/doi/10.1126/science.aat7615. Cell-type specific latent factor 4 loadings are published in https://www.nature.com/articles/s41586-024-07109-5.

## Acknowledgements

R.L.S is a PhD candidate in the NIH Oxford-Cambridge Scholars Program. All research from the Department of Psychiatry at the University of Cambridge is made possible by the NIHR Cambridge Biomedical Research Centre and the NIHR East of England Applied Research Centre. The views expressed are those of the author(s) and not necessarily those of the NHS, the NIHR or the Department of Health. This work received computational support from the NIP HPC Biowulf cluster (http://hpc.nih.gov) and from the mental health theme of the National Institute of Health Research (NIHR) Cambridge Biomedical Research Center. Library preparation and RNA sequencing was performed at the NIH Sequencing Center. Thank you to families of the deceased for tissue donations and to the Offices of the Medical Examiner of the District of Columbia, Central and Northern Virginia for referrals and brain extraction.

## Author Contributions

Conceptualization, R.L.S., A.M. A.R., P.E.V., F.J.M.; methodology, R.L.S., A.M., N.A.; software, R.L.S., A.M..; formal analysis, R.L.S., A.M.; data curation, N.A., P.K.A., S.M., F.J.M.; writing – original draft, R.L.S., A.R., P.E.V., F.J.M.; writing – review and editing, R.L.S., A.M., N.A., P.K.A., S.M., A.R., P.E.V., F.J.M.; visualization, R.L.S.; supervision, A.R., P.E.V., F.J.M.; project administration, N.A., P.K.A., S.M., F.J.M.

