## Supporting Information for "A neuro-immune axis of transcriptomic dysregulation within the subgenual anterior cingulate cortex in schizophrenia"

##### This PDF file includes

Supporting text
Figs S1 to S15
SI References

##### Other supporting materials for this manuscript include the following

Tables S1 to S10

### Supporting text

#### Methods

##### *WGCNA module assignment*

The WGCNA modules used in the current work were selected after careful consideration across parameters used in the WGCNA R package [1]. First, we selected the soft threshold power by comparing scale-free topology model fit across a range of potential powers (1 through 16, inclusive). We constructed a signed network to retain information about the directionality of node correlation. We selected a soft threshold power of 3 because it was the lowest power exceeding an $R^2$ threshold of 0.8 while maintaining relatively high median connectivity (around 100 or more; **Fig S4A**). Although we considered a soft threshold power of 2, it did not yield modules with robust biological enrichment.

After constructing the network using a soft threshold power of 3, we examined module assignments across varying minimum module sizes and cut heights (**Fig S4B**). Minimum module sizes ranged from 30 to 50 in increments of 5 (30, 35, 40 shown in **Fig S4B**), while cut heights ranged from 0.90 to 0.98 in increments of 0.003 (0.95 to 0.98 shown in **Fig S4B**). For each parameter combination, we evaluated module number, size, and biological enrichment, aiming to capture distinct biological processes without excessive fragmentation. Particular attention was given to the size of the gray module, ensuring it remained below ~10,000 genes to avoid conflating potentially significant signals.

##### *Topic modeling*

Topic modeling is a text mining method for unsupervised classification of text that facilitates understanding of the semantic organization of functional enrichments (**Fig S5B**). The algorithm identifies natural groups of co-occurring words in significant GO terms, or “topics”. The algorithm can then quantify the mixture of words associated within each topic, while also determining the mixture of topics that describes each grouping (in this case – reference module). Here, Frequently in and Exclusively (FREX) words are used to characterize each topic. The strength of association of each word with each topic is described by the parameter beta. The strength of association of each gene list module with each topic is described by the parameter gamma, which is the estimated proportion of words from the GO terms in that module that are generated from the respective topic. Thus, each module is summarized by general biological function across all pathway results. Number of topics (K = 5) was chosen based on optimal exclusivity and semantic coherence, biological knowledge, strength of association between gene lists and topics.

##### *WGCNA module eigengene analysis*

To assess the relationship between WGCNA modules and psychiatric disorders, we used the canonical module eigengene approach [2]. For each module, we ran principal components analysis (PCA) on the gene-by-sample expression matrix. We extracted PC1 as the module eigengene. We then ran a linear regression model for each module, with module eigengene as the outcome variable, and psychiatric disorder and the first 8 drug MCA dimensions (eigengene ~ diagnosis + MC1 + MC2 + MC3 + MC4 + MC5 + MC6 + MC7 + MC8). A module-covariate association was considered significant if the term $P < 0.05$.

##### *Published neuropsychiatric risk gene lists*

Gene-disorder associations across methods (GRCCA, DGE, WGCNA) were benchmarked against the psychiatric genetics literature using identified risk genes across disorders. Risk gene lists were based on colocalization of genes with genome-wide significant variants as follows:

- SCZ broad set (common variants; 95% credible set, k<=3.5; N=629) [3]
- SCZ prioritized (common variants with convergent SMR/TWAS results; N=120) [3]
- SCZ genes implicated through whole exome sequencing meta-analysis (rare variants; SCHEMA; N=10) [4]
- BD common variant-associated genes identified using MAGMA (N=162) [5]
- MDD common variant-associated genes identified using MAGMA (N=339) [6]
- Autism Spectrum Disorder (ASD) common variant-associated genes identified using H-MAGMA (N=567) [7]

#### Results

##### *Multiple correspondence analysis summarizes toxicology data across post-mortem samples*

To quantify the extent to which medicine and recreational drug confounders contribute to gene expression data, toxicology data were included in the GRCCA covariate (Y) matrix. Seventeen distinct drugs were detected in at least one postmortem sample and recorded as a binary variable (detected (1) or not detected (0); **Fig S3A**). Multiple correspondence analysis was run on drug covariate data across samples to reduce dimensionality in GRCCA without losing information. The first eight dimensions were selected for subsequent analyses and explained 76.6% of the variation in the data (**Fig S3B**). Of the eight dimensions we identified, the first dimension identified detection of any compound (**Fig S3C**; **Fig S3D**). Dimension two was driven by anti-anxiety medication (benzodiazepines and sedatives), dimension 3 by mood stabilizers and anti-epileptic medication, dimension 4 by antihistamines and ‘other’ psychotropic drugs, 5 by antihistamines and cannabinoids, dimension 6 by alcohol in the positive direction and cannabinoids in the negative, 7 by major stimulants including cocaine, and 8 by cannabinoids (**Fig S3C**).

##### *Weighted gene co-expression analysis*

WGCNA identified 23 modules in the expression data of 18,667 genes (**Fig S5A**; **Table S7A**). GO functional enrichment revealed that every module was significantly (FDR or *P* < 0.05) related to various known biological functions, including: synaptic signaling, mitochondrial function, transcription and translation, cellular development and differentiation, transport and biosynthetic processes, and immune function (**Table S7B**; **Fig S5B**). Cell-type enrichment analysis demonstrated that each module, excepting geneM7 and geneM14, was significantly (hypergeometric *P* or FDR < 0.05) enriched for at least one cell type, including astrocytes, endothelial cells, excitatory and inhibitory neurons, microglia, oligodendrocytes, and oligodendrocyte precursor cells (**Fig S5C**). All modules displayed dynamic patterns of expression across normative development (**Fig S5D**).

##### *Module eigengene analysis links gene co-expression to SCZ*

Gene-level modules were associated with psychiatric disorders using the canonical module eigengene approach as described in **Methods**. Modules geneM2, geneM9, and geneM15 were all associated with SCZ (**Fig S12A**; **Table S8**). These modules were enriched for cell transportation and development, vesicle binding and synaptic signaling, and voltage-gated ion channel activity, respectively (**Fig S5**; **Fig S12B**). However, none of these modules were enriched for SCZ risk genes, as determined by hypergeometric testing (**Fig S13**).

##### *Co-expression patterns of significant GRCCA genes (module-level GRCCA results)*

Hypergeometric testing of each module with the significant GRCCA gene subset (**Fig S8**) in that module revealed geneM1, geneM4, and geneM14 as overrepresented in the GRCCA results (FDR < 0.01; **Fig S14A**). Significant GRCCA genes in geneM1 and geneM4 have generally positive structure correlations (indicating up-regulation in SCZ), and significant GRCCA genes in geneM14 generally have negative structure correlations (indicating down-regulation in SCZ; **Fig S14B**). Modules geneM1 and geneM4 are enriched for vesicle transport, ion channel activity, and synaptic signaling (**Fig S5**; **Fig S14C**), while genes in module geneM14 are enriched for metabolic activity and mitochondrial respiration (**Fig S14C**). Notably, SCZ risk genes are also enriched in modules geneM1 and geneM4 (**Fig S13**).

##### *Exclusion of the grouping vector (RCCA) leads to weaker risk gene enrichment*

To assess the extent to which the inclusion of the grouping vector influences the multivariate (GRCCA) results, we also ran RCCA on the same data. The feature-level regularization parameter (lambda) was set to 1 - 1/(n features) (same as GRCCA), but the grouping vector was excluded (i.e., underlying data structure was not considered in this model). The RCCA model was optimized at 40% variance ($P$ = 0.002; **Table S9A**), indicating that less variance in gene expression data could be included without overfitting. This is unsurprising as the grouping vector in GRCCA provides additional regularization. The RCCA covariate results were almost identical to GRCCA covariate results (**Table S9B**; **Fig S15A**), with SCZ RCCA $r_x$ = 0.917 vs SCZ GRCCA $r_x$ = 0.920. Gene structure correlations were also very highly correlated (**Fig S15B**; **Table S9C**), and cell-type and GO pathway enrichments were similar to GRCCA enrichments (**Table S10A** and **Table S10B**). Using the $|Z| > 2$ and FDR < 0.05 criteria, 2026 genes were significant in the RCCA analysis, 855 of which were also significant in GRCCA (**Fig S15C**; hypergeometric $P$ < 0.001). Together, these results demonstrate that the GRCCA grouping vector did not drastically affect the multivariate results. However, it did appear to fine tune them, as the SCZ risk gene enrichment was stronger in GRCCA than RCCA, especially for the prioritized variant list (**Fig S15D**; **Table S10C**).

#### Supporting Figures S1 to S14

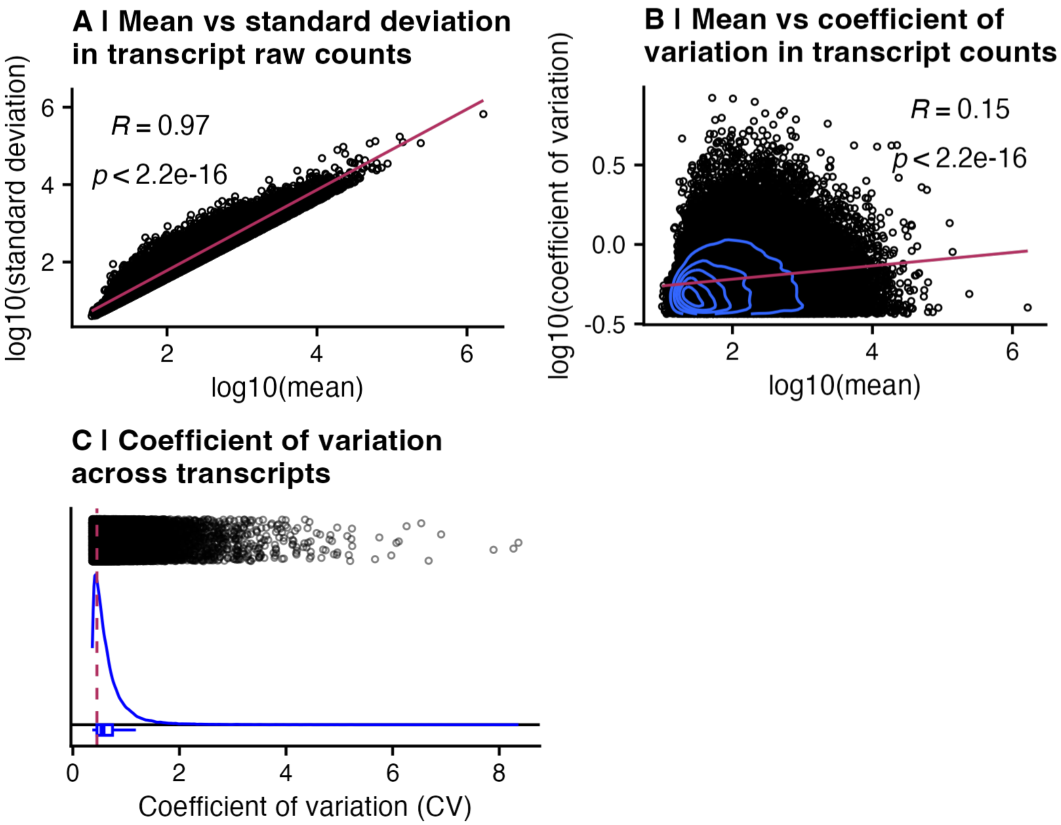


**Figure S1. Transcript filtering criteria for inclusion in analysis. (A)** The relationship between transcript mean expression (x-axis) and standard deviation of expression (y-axis). In general, genes with higher expression also have higher variance in expression levels. Thus, excluding transcripts solely based on low variances would selectively remove transcripts with low expression levels. **(B)** The relationship between mean transcript expression (x-axis) and transcript coefficient of variation (standard deviation / mean; y-axis). Though the two are less correlated than mean and standard deviation. **(C)** The coefficient of variation (CV = standard deviation / mean; x-axis) filtering criteria. Transcripts in the bottom quantile (25%) of CV (i.e., low variance relative to mean) were excluded from downstream analysis due to lack of significant variation across samples.

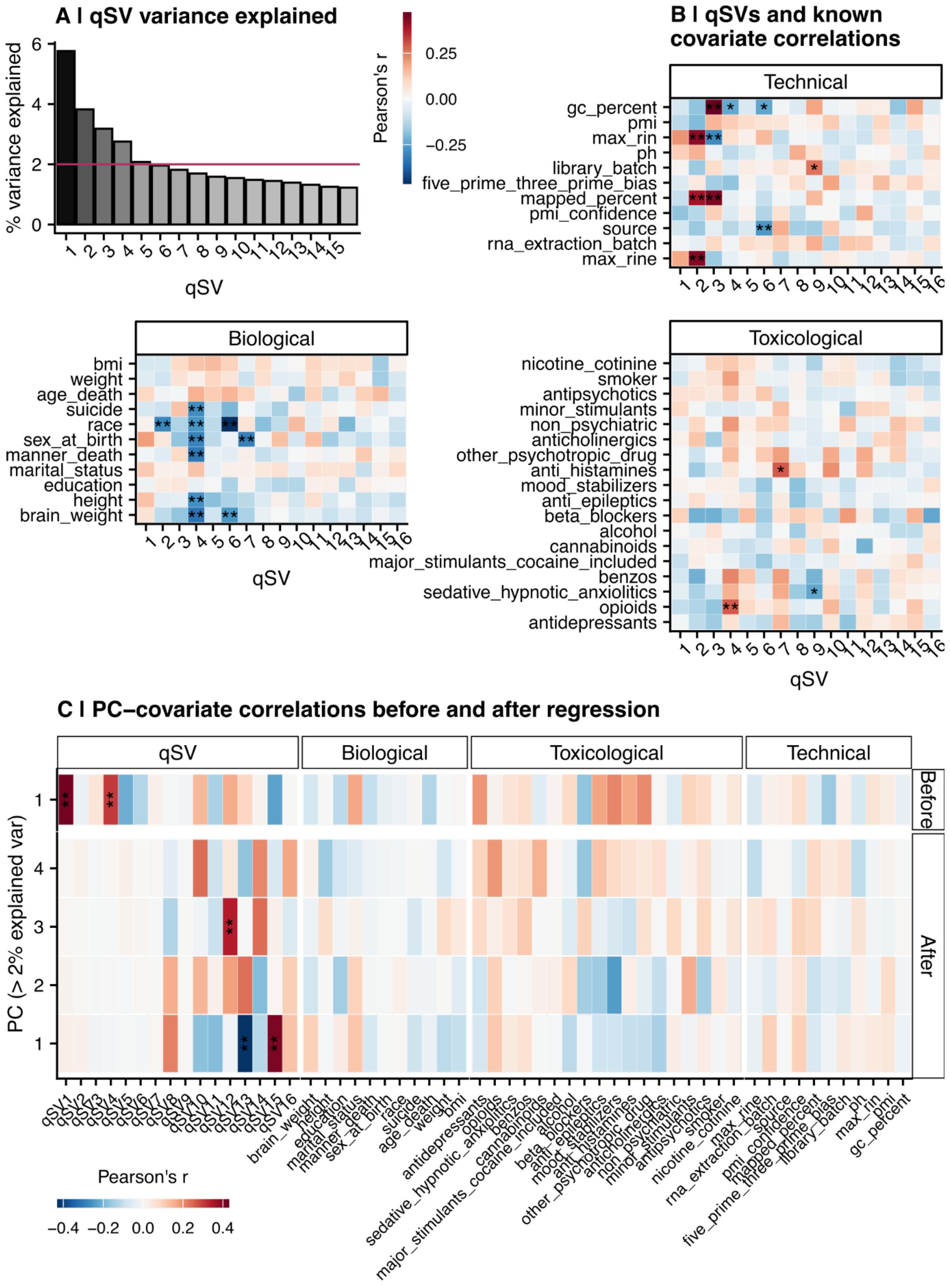
A I qSV variance explained
% variance explained
qSV
Pearson's r
0.25
0.00
−0.25
B I qSVs and known covariate correlations
Technical
gc_percent
pmi
max_rin
ph
library_batch
five_prime_three_prime_bias
mapped_percent
pmi_confidence
source
rna_extraction_batch
max_rine
Biological
bmi
weight
age_death
suicide
race
sex_at_birth
manner_death
marital_status
education
height
brain_weight
Toxicological
nicotine_cotinine
smoker
antipsychotics
minor_stimulants
non_psychiatric
anticholinergics
other_psychotropic_drug
anti_histamines
mood_stabilizers
anti_epileptics
beta_blockers
alcohol
cannabinoids
major_stimulants_cocaine_included
benzos
sedative_hypnotic_anxiolitics
opioids
antidepressants
C I PC−covariate correlations before and after regression
qSV
Biological
Toxicological
Technical
PC (> 2% explained var)
Before
After
Pearson's r
−0.4 −0.2 0.0 0.2 0.4

**Figure S2. Covariate regression from transformed raw count data. (A)** Variance in transcript degradation data explained by qSVs. Each qSV is shown on the x-axis, while the proportion of variance explained iss shown on the y-axis and indicated by bar color. The maroon dashed line shows the threshold for qSV inclusion in the regression model (proportion of variance explained >= 0.02). **(B)** Correlations between qSV and known covariates (biological, technical, and toxicological; **Table S1**). The x-axis shows each qSV, while the y-axis includes all known covariates. Tiles are colored by the Pearson correlation (where red indicates $r > 0$ and blue indicates $r < 0$). Significance is indicated by number of asterisks (FDR < 0.05 ~ *, FDR < 0.01 ~ 0.01). **(C)** The Pearson correlation between each significant PC (proportion of variance explained >= 0.02) in the gene expression data (x-axis) and known covariates (**Table S1**) and all qSVs (y-axis) before (top) and after (bottom) covariate regression. Tiles are colored by the Pearson correlation (where red indicates $r > 0$ and blue indicates $r < 0$). Significance is indicated by number of asterisks (FDR < 0.05 ~ *, FDR < 0.01 ~ 0.01). Note that qSVs 12, 13, and 15 were not regressed since they did not account for at least 2% of the variance or correlate with a known covariate.

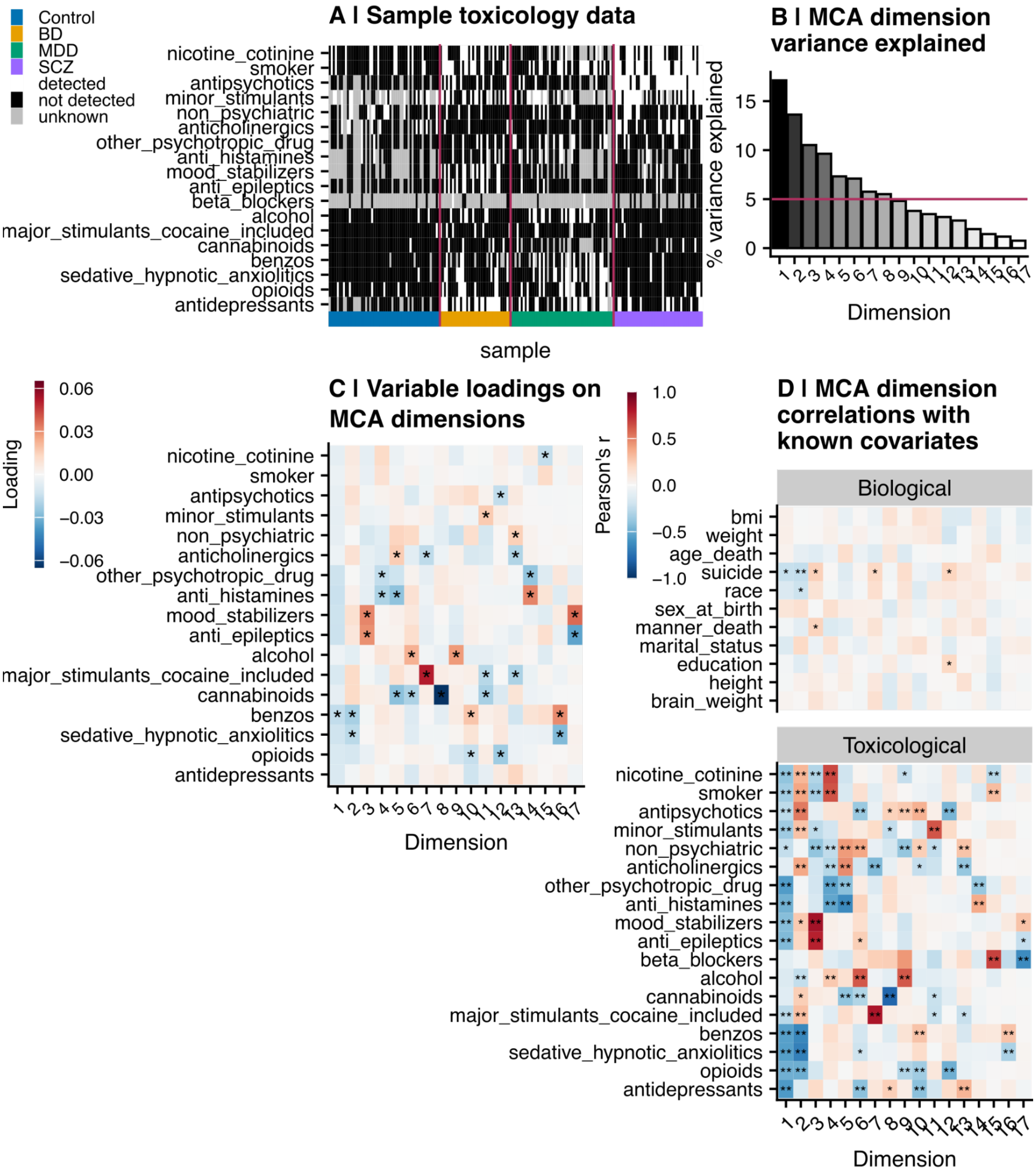


**Figure S3. Toxicology multiple correspondence analysis (MCA) results. (A)** Matrix representation of toxicology reports across 185 post-mortem samples included in this study. The y-axis indicates each medication or recreational drug reported or detected by toxicology. The x-axis shows samples, ordered by diagnostic group. Each tile is colored by drug use coded as a binary variable (black = drug not reported or detected in toxicology, white = drug reported or detected, gray = not reported/known). The vertical maroon lines separate each diagnostic group. **(B)** Variance explained of each dimension in the MCA of 17 known compounds. Each dimension is shown on the x-axis, and its associated percentage of variance explained is shown on the y-axis and indicated by bar color. The horizontal maroon line indicates the threshold for inclusion in subsequent models (proportion of variance explained > 0.02). **(C)** Compound loadings on each MCA dimension. Loadings with absolute value greater than 0.015 are indicated by an asterisk. **(D)** Pearson correlation of each MCA dimension with each known covariate. Panel legend is the same as **Fig S2B**, with MCA dimension on the x-axis (rather than principal component).

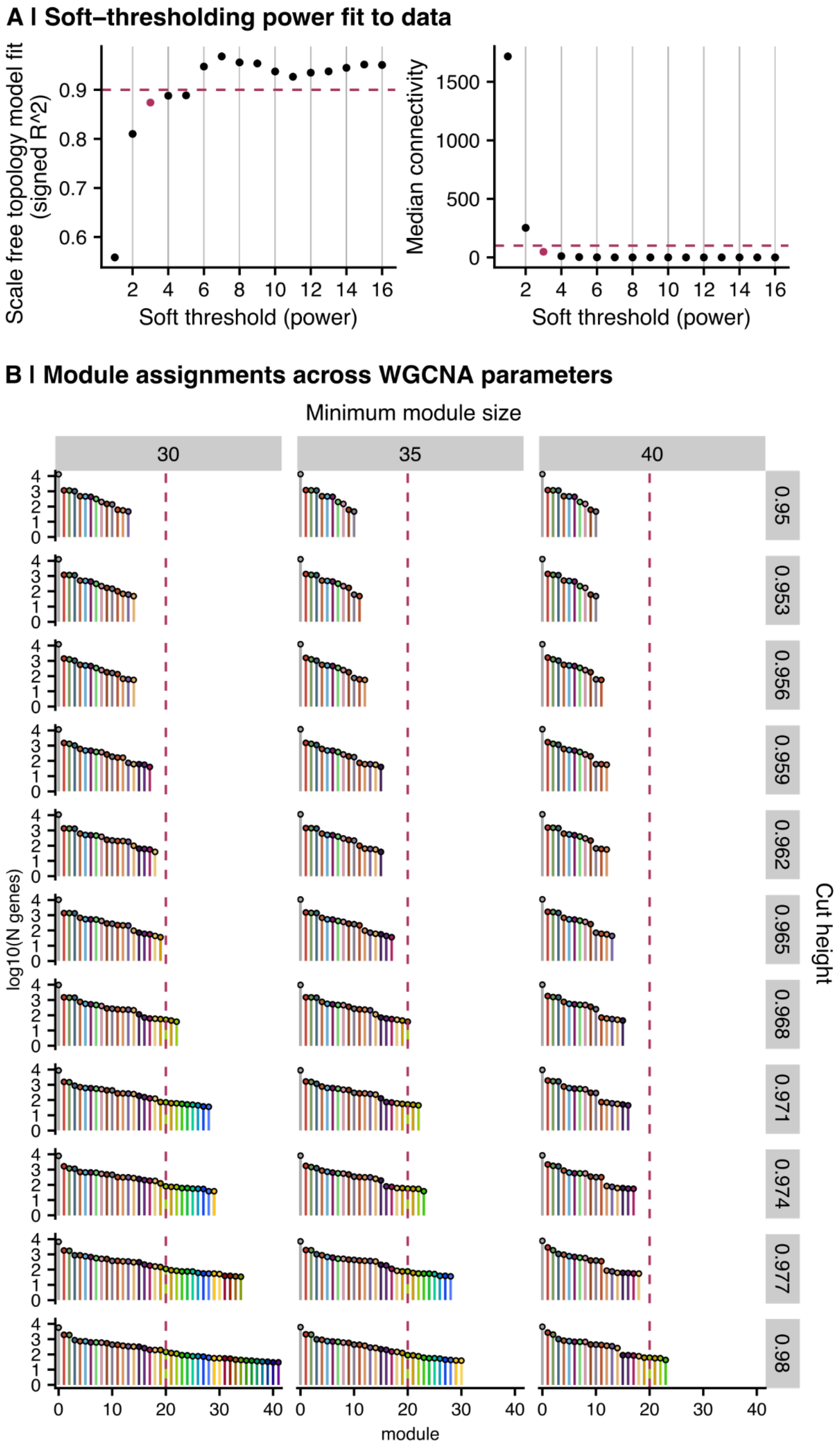
A | Soft–thresholding power fit to data
Scale free topology model fit (signed R^2)
Soft threshold (power)
Median connectivity
Soft threshold (power)
B | Module assignments across WGCNA parameters
Minimum module size
30
35
40
0.95
0.953
0.956
0.959
0.962
0.965
0.968
0.971
0.974
0.977
0.98
Cut height
log10(N genes)
module

**Figure S4. Selecting WGCNA parameters for module assignment. (A)** Soft power threshold selection. Left: Model fit ($R^2$; y-axis) across soft threshold powers (x-axis). Package authors recommend a model fit around 0.9 (represented by maroon dashed line). Right: Median connectivity (y-axis) across soft-threshold powers (x-axis). Package authors recommend median connectivity around 100 or higher (represented by maroon dashed line). The selected soft threshold power of three is highlighted in maroon. **(B)** Minimum module size and cut height selection at soft threshold power = 3. The module number is shown on the x-axis, and the log10-transformed gene count per module is represented by the y-axis. Columns are faceted by minimum module size; rows are faceted by tree cut heights. Module number 20 is highlighted by the maroon dashed line as the approximate target number of modules.

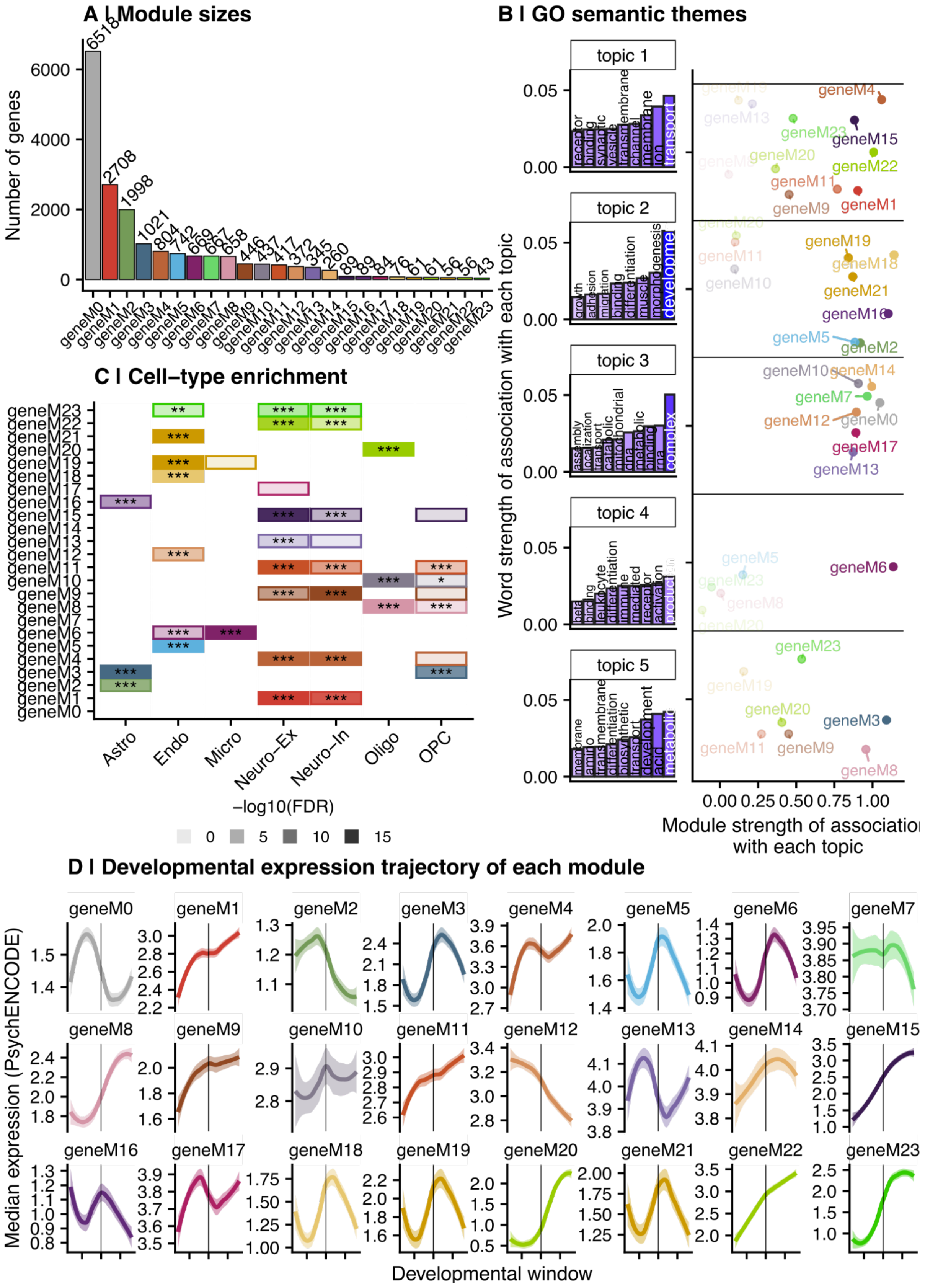
A I Module sizes
Number of genes
B I GO semantic themes
topic 1
topic 2
topic 3
topic 4
topic 5
Word strength of association with each topic
Module strength of association with each topic
C I Cell–type enrichment
Astro
Endo
Micro
Neuro–Ex
Neuro–In
Oligo
OPC
−log10(FDR)
D I Developmental expression trajectory of each module
Median expression (PsychENCODE)
Developmental window

**Figure S5. WGCNA identifies 23 biological robust gene co-expression modules. (A)** Number of genes in each module. The WGCNA module is shown on the x-axis and represented by bar color. The height of the bar indicates the number of genes assigned to the respective module. **(B)** Functional enrichment themes across modules as determined by topic modeling. **Left**: The strength of association of the top 10 words with each topic. Each bar represents a FREX word (frequently and exclusively associated with the given topic), while the y-axis, bar color, and word size all indicate the strength of association of that word with the respective topic, as determined by the topic model algorithm. **Right**: The strength of association of each module with each topic. The module strength of association is represented on the x-axis, where 0 indicates no association and 1 suggests the module aligns perfectly with the respective topic. The y-axis represents the topic, faceted to align with the left panel. Each module is represented by a point and labeled and colored accordingly. **(C)** Cell-type enrichment of WGCNA modules by hypergeometric test. The x-axis indicates each cell type, as defined by [8]. The y-axis shows each WGCNA module. Tiles represent a significant hypergeometric overlap between the respective cell type and WGCNA module. Tiles are colored by module, and tile transparency corresponds to -log10(FDR). * = FDR < 0.05; ** = FDR < 0.01; *** = FDR < 0.001. **(D)** Mean expression pattern of genes in each module across development. The x-axis indicates developmental window, defined by [9], where the vertical line separates pre- (left) and post- (right) natal windows. The y-axis shows the expression value across frontal cortex samples, as reported in [9]. Each facet represents a module, where the line shows the loess curve of mean expression across genes within the respective module. Lines are colored by module.

**A l RCCA setup**

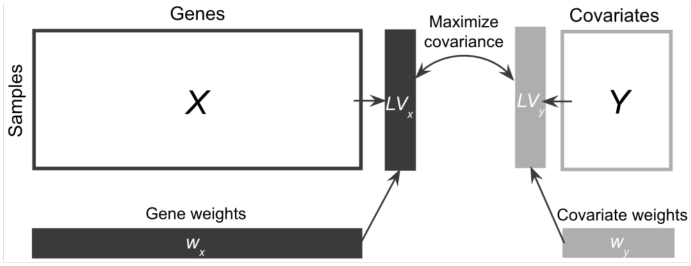


**B l GRCCA setup**

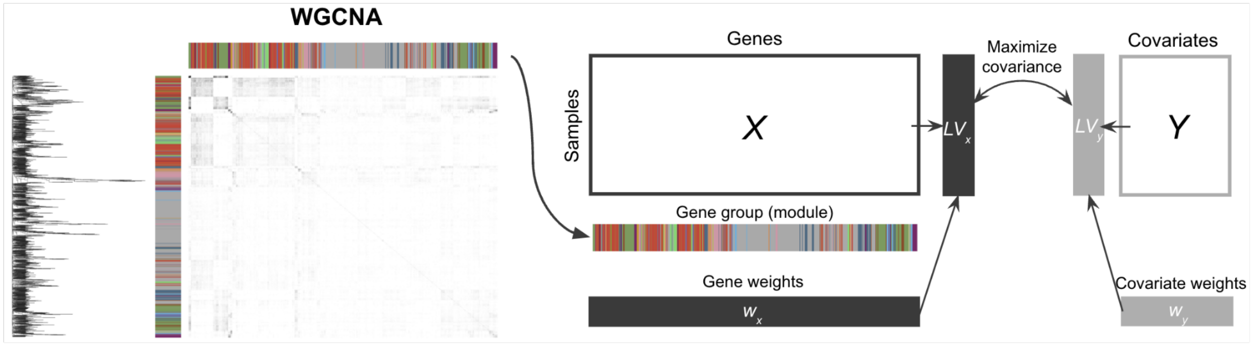


**Figure S6. Schematic of analysis design for CCA analyses. (A)** Regularized CCA (RCCA) analysis design to identify the maximized linear association between genes and covariates (including diagnosis and medications/recreational drugs). Briefly, the *X* matrix is gene expression across samples data, while the *Y* matrix is covariate data across samples (with diagnostic group represented as a dummy matrix). The RCCA algorithm determines a vector of gene weights ($w_x$) and separately, a vector of covariate weights ($w_y$) that, when multiplied by their respective data to calculate the latent variables (i.e., $LV_x = X \cdot w_x$ and $LV_y = Y \cdot w_y$), maximally covary. Feature loadings, or in the case of CCA, structure correlations, are calculated as the correlation between the data and the latent variable (i.e., cor($X$, $LV_x$) and cor($Y$, $LV_y$)). **(B)** Group RCCA (GRCCA) analysis schematic. The inputs, outputs, and underlying algorithm of GRCCA is largely the same as RCCA. The primary difference is that while RCCA assumes feature independence, GRCCA accounts for underlying structure in the data by regularizing at a group level, in addition to the feature level. Here, feature (gene) group is determined by WGCNA module, which is given to the algorithm as a 1 x (n features) vector. The GRCCA outputs are the same as RCCA (meaning, there is no group-specific output).

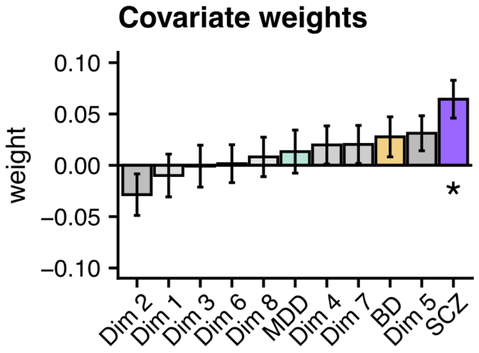


**Figure S7. Gene-level GRCCA covariate weights.** Covariate (Y matrix) weights ($w_y$) as determined by the GRCCA algorithm. Note that these are distinct from (but related to) the structure correlations (cor($Y$, $LV_y$ = $Y \cdot w_y$)) reported in the main text. The x-axis shows each covariate, and the y-axis shows their respective weights (+/- standard deviation across 1000 bootstraps). * = $FDR_r$ < 0.05 & |$Z$| > 2.

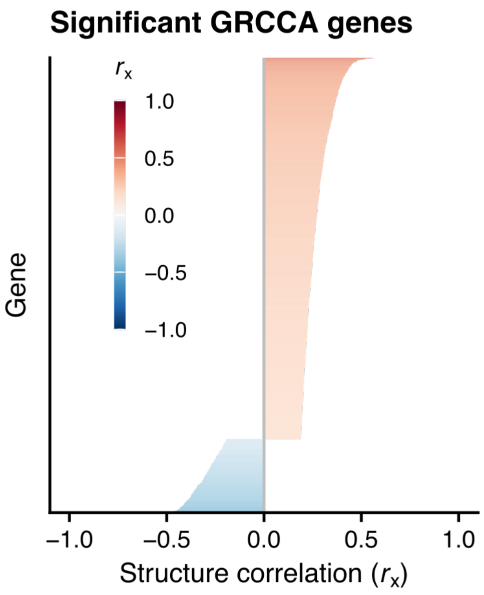


**Figure S8. Vector of all significant GRCCA genes.** Representation of all genes considered significant at $FDR_r < 0.05$ & $|Z| > 2$ per the GRCCA algorithm (N = 1211). The x-axis shows structure correlation ($r_x$) while each column on the y-axis represents a gene (colored by $r_x$).

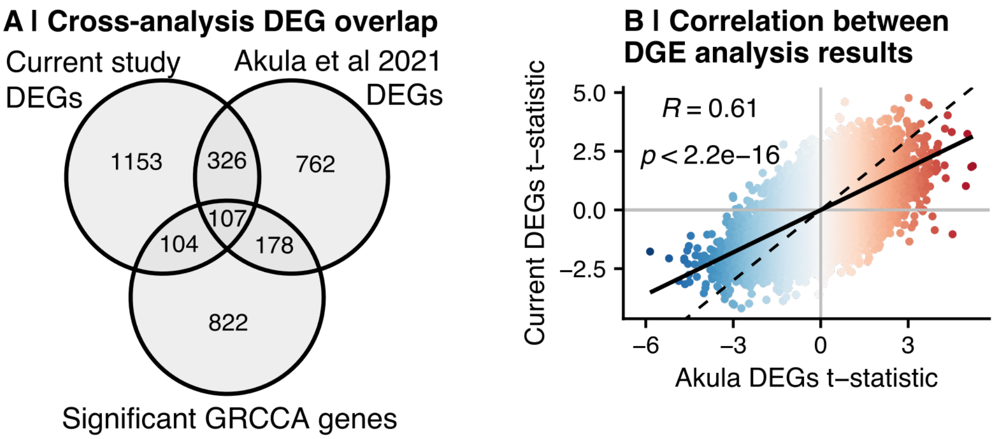


**Figure S9. Comparison between current study DEGs and Akula et al 2021 DEGs. (A)** A venn diagram showing the overlap among significant DEGs from the current study (N = 1690), significant DEGs from [10] (N = 1373), and significant GRCCA genes (N = 1211). The primary difference between the two DGE analyses is the raw count preprocessing pipeline. Pairwise hypergeometric test results are as follows: Current DEGs vs Akula DEGs: N = 433; $P < 0.001$; Current DEGs vs GRCCA genes: N = 211, $P < 0.001$; Akula DEGs vs GRCCA genes: N = 285, $P < 0.001$. **(B)** A scatter plot showing the correlation between [10] log2(fold change) effect size (x-axis) and the normalized log2(fold change) effect size (y-axis) across genes in the current study. Each point represents a gene, and is colored by the log2(fold change) in [10], where red indicates an increase in expression in SCZ compared to controls, and blue indicates a decrease. The solid line indicates the line of best fit ($r$ = 0.61), while the dashed line represents the identity line (y = x).

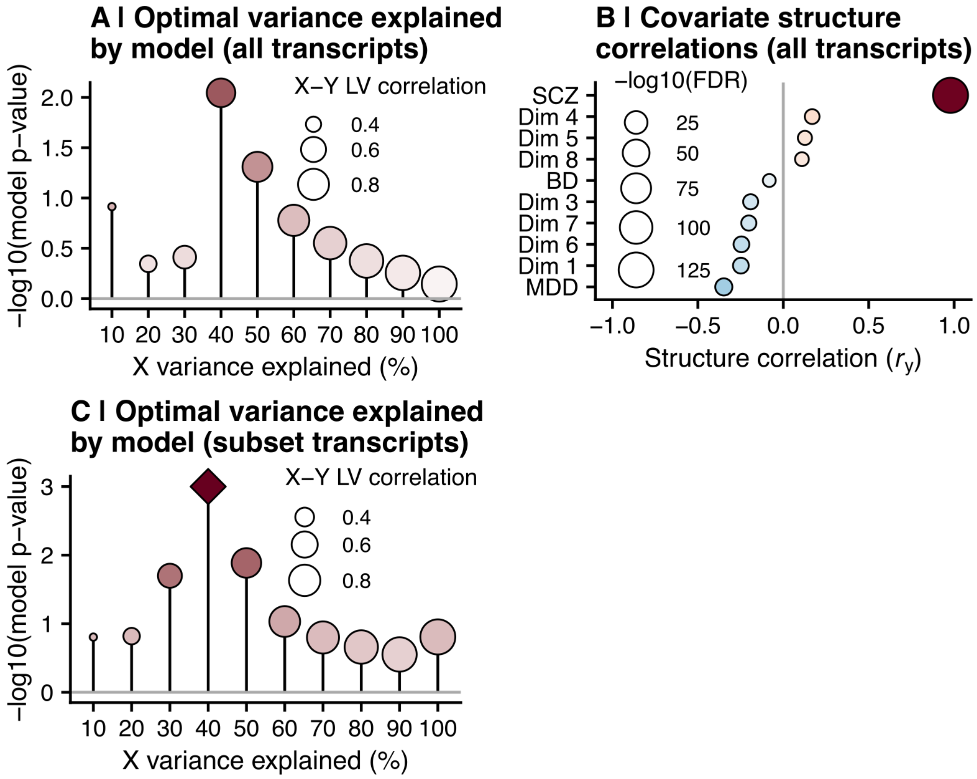


**Figure S10. Transcript-level GRCCA model results. (A)** GRCCA model *P* values for the transcript-level analysis that included all transcript derivatives that met the filtering thresholds (**Supplement**; N = 54,302). Figure legend is the same as **Fig 2A**. No model met the significance threshold of *P* = 0.001 (the most significant model was optimized at 40% variance in the expression data, *P* = 0.009). **(B)** Covariate structure correlations for the GRCCA model that included all transcript (N = 54,302; optimized at variance explained = 40%). SCZ was the covariate most strongly associated with the latent variable ($r_y$ = 0.979; FDR < 0.001; $Z$ = 1.39). **(C)** GRCCA model *P* values for the transcript-level analysis reported in the main text (**Results**; including only transcript derivatives of 1. common variant-associated genes of any psychiatric disorder, or 2. genes significantly associated with the latent variable in the gene-level GRCCA (at FDR < 0.05); N = 12,986). Figure legend is the same as **Fig 2A** and **Fig S10A**.

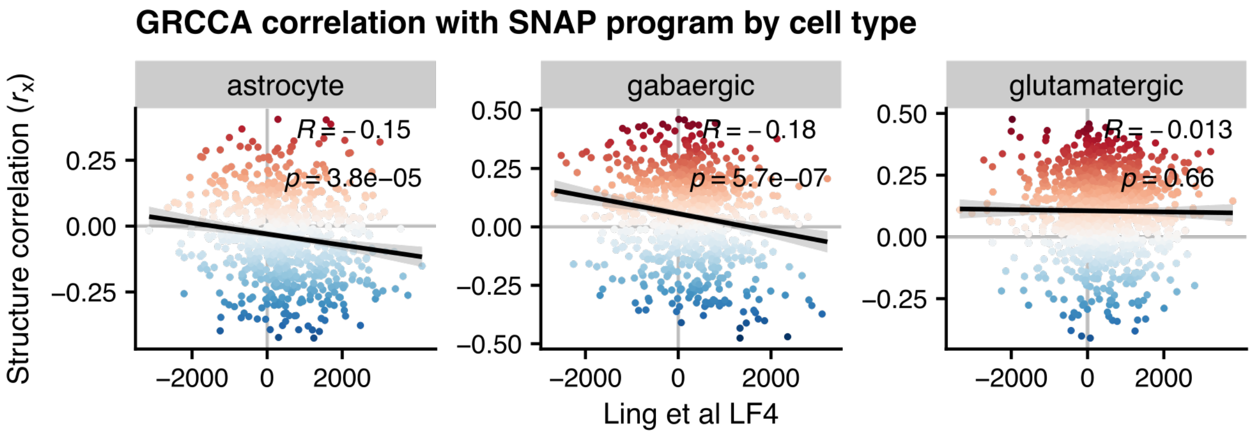


**Figure S11. GRCCA results are related to expression gradients in the single-cell schizophrenia literature.** The cell-type specific relationship between latent factor 4 (LF4) from Ling et al (2024) [11] and GRCCA structure correlation. Each point represents a gene specific to a cell type in both Ling et al (2024) [11] and the current study. The x-axis represents its loading on LF4, while the y-axis represents its loading (structure correlation) on the GRCCA latent variable. Points are colored by GRCCA structure correlation. Ling and colleagues [11] demonstrated that LF4 decreases in schizophrenia and in aging, while GRCCA structure correlation is positively associated with schizophrenia; thus, a negative correlation between the two indicates directionality alignment. The lack of correspondence between gene structure correlation and LF4 loading in glutamatergic cells likely reflects regional cell-type heterogeneity (i.e., inhibitory cells are more abundant in the sgACC, and excitatory neuron subclusters demonstrate considerable diversity across brain regions) [12].

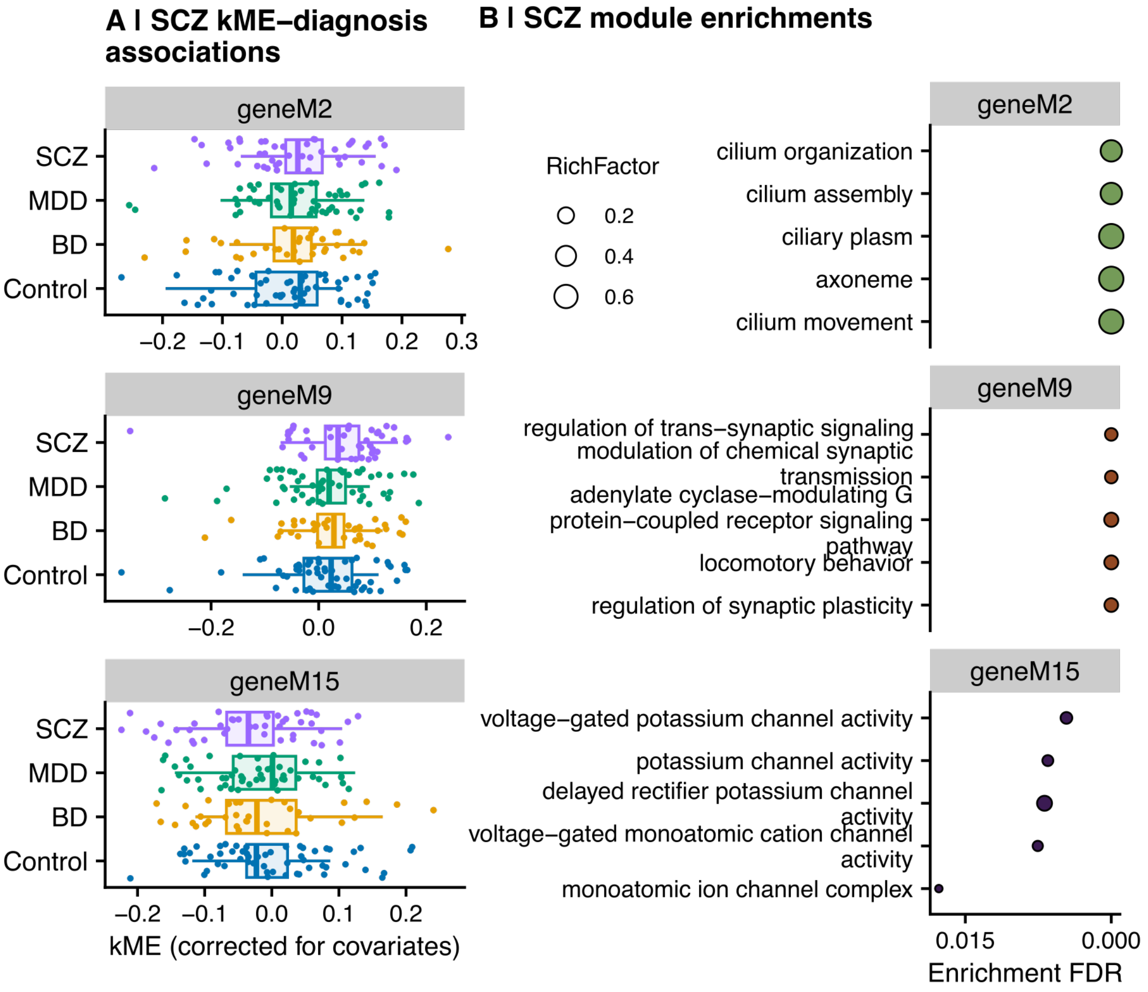


**Figure S12. SCZ-WGCNA module associations as determined by canonical module eigengene analysis. (A)** The distribution of module eigengene by diagnostic group in the three modules (geneM2, geneM9, geneM15) that were significantly associated with SCZ. For each module, a linear model of eigengene (kME) on diagnosis was run, with MCA drug dimensions 1-8 as covariates (to maintain model consistency across methods). All three modules demonstrated a significant difference in kME distribution across SCZ samples at $p < 0.05$, though none of the associations were significant after FDR correction. The x-axis represents residualized kME, while the y-axis indicates diagnostic group. Each point represents a sample and is colored by its associated diagnosis, while the boxplot shows the overall distribution for the group. **(B)** GO biological process pathway enrichments for each module. The top five pathways by FDR for each module are shown on the y-axis, while the x-axis represents the FDR-corrected p-value. Each point is sized by the pathway rich factor and colored by module.

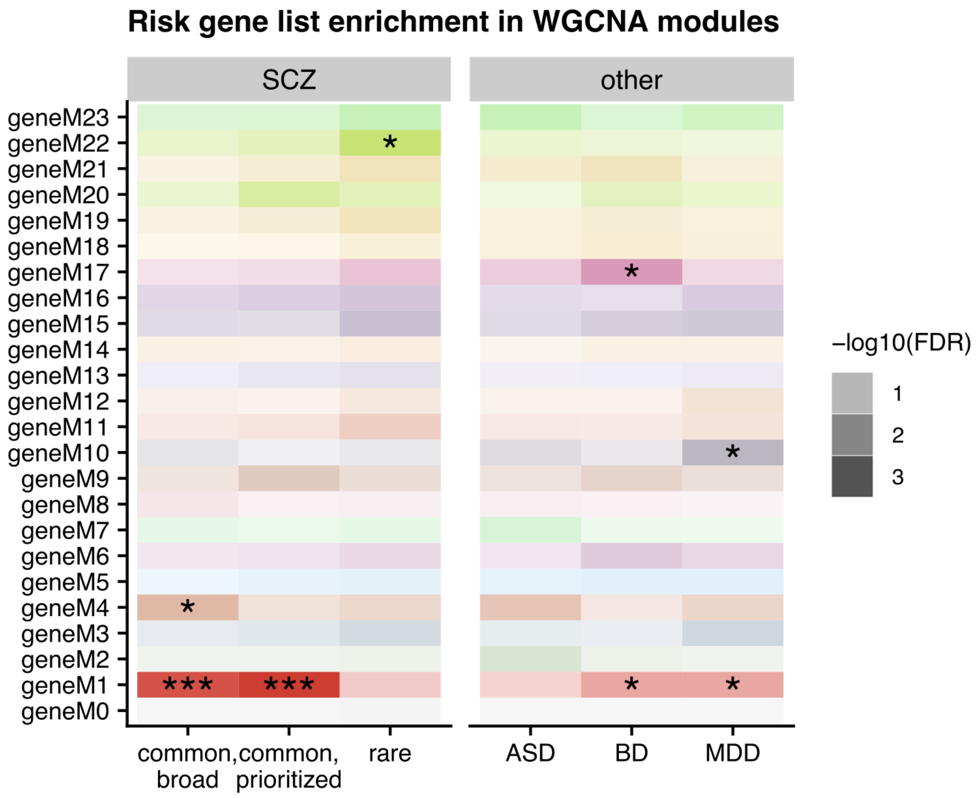


**Figure S13. Risk gene enrichment in WGCNA modules.** Hypergeometric overlap between a risk gene list associated with psychiatric disorders (x-axis) and WGCNA module (y-axis). The tile color also represents the WGCNA module, while the opacity indicates overlap significance (darker = smaller FDR). * = FDR < 0.05; ** = FDR < 0.01; *** = FDR < 0.001.

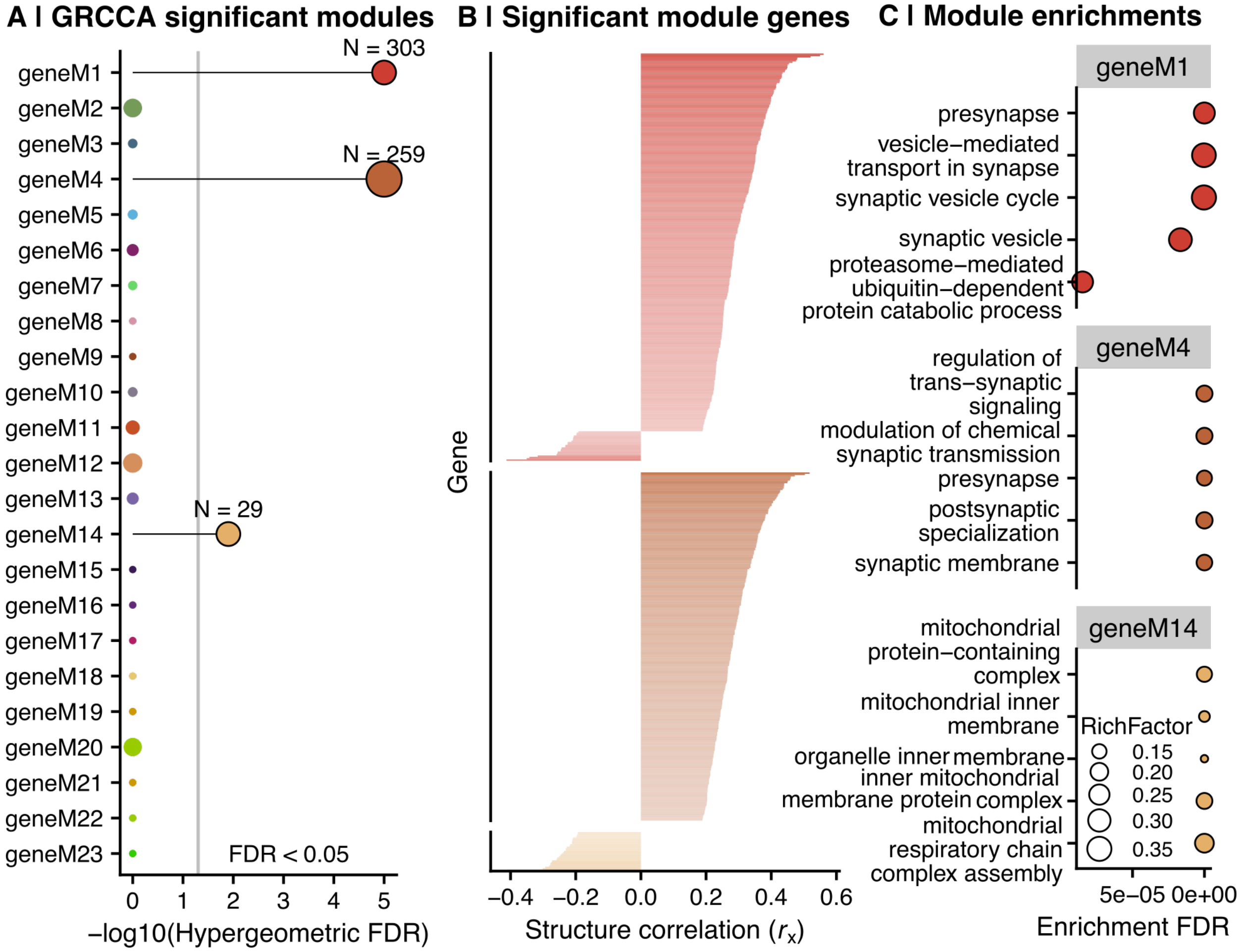


**Figure S14. WGCNA modules that are overrepresented in the GRCCA gene list. (A)** Hypergeometric overlap of WGCNA modules and significant GRCCA genes. The y-axis shows each WGCNA module and the x-axis represents its -log10(FDR-corrected hypergeometric p-value). The horizontal line shows FDR = 0.01. Points are colored by module (per **Fig S4**) and those that were significant at FDR < 0.01 are outlined in black. The size of the point indicates the number of genes in the module-GRCCA intersect. **(B)** The structure correlations of significant genes in the overrepresented modules (FDR < 0.01). Each horizontal bar is a gene (ordered on the y-axis by increasing structure correlation within module), and the x-axis shows its structure correlation. **(C)** GO biological process pathway enrichments for each module. Panel legend is the same as **Fig S12B**.

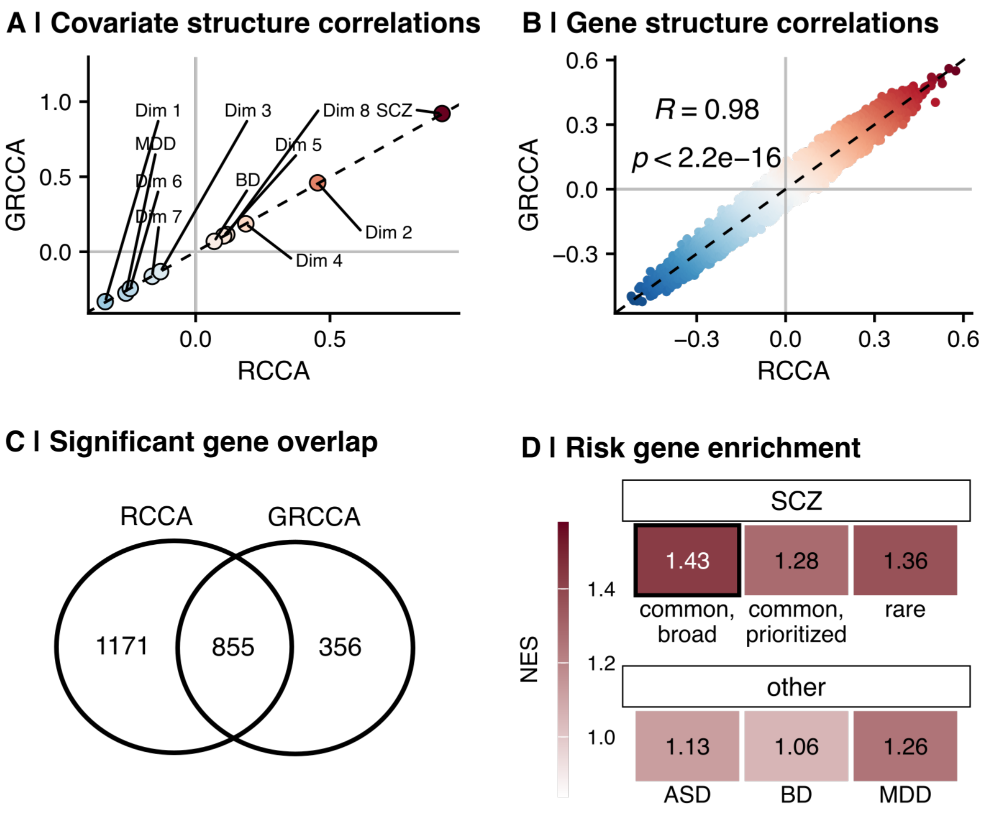


**Figure S15. A direct comparison of RCCA and GRCCA results. (A)** Correlation between RCCA covariate structure correlations (x-axis) and GRCCA covariate structure correlations (y-axis) ($R$ = 1, $P$ < 0.001). Each point represents a covariate (labeled) and is colored by the sum of the GRCCA and RCCA structure correlations. The dashed black line indicates y = x. **(B)** Correlation between RCCA gene structure correlations (x-axis) and GRCCA gene structure correlations (y-axis) ($R$ = 0.98, $P$ < 0.001). Each point represents a gene and is colored by the sum of the GRCCA and RCCA structure correlations. The dashed black line indicates y = x. **(C)** A venn diagram representing the intersection of significant genes as determined by RCCA (teal) and GRCCA (salmon). The text indicates the number of genes in each category; circles are not drawn to scale. Hypergeometric $P$ < 0.001. **(D)** Enrichment of RCCA results for psychiatric disorder risk genes. Figure legend is the same as **Fig4A**.
